# Characterisation of the new microalgal protein xATPA related to the F-type ATP synthase α subunit, from the ecosystem to the molecule

**DOI:** 10.64898/2026.07.08.737214

**Authors:** Mathias Penot-Raquin, Anna M. G. Novák Vanclová, Victoria Powell, Yann Corbeau, Chaima Younes, Maxime Eugène, Tahar Bouceba, Cédric Pionneau, Viviane de Almeida Bastos, Mathilde Garcia, Chris Bowler, Richard G. Dorrell

**Affiliations:** Institut de biologie de l’École normale supérieure (IBENS), École normale supérieure, CNRS, INSERM, Université PSL, 75005 Paris, France; Sorbonne Université, CNRS, Computational, Quantitative and Synthetic Biology, CQSB, F-75005, Paris, France; Roscoff Culture Collection, Sorbonne Université, Station Biologique de Roscoff, Roscoff 29680, France; Sorbonne University, CNRS, Institut de Biologie Paris-Seine (IBPS), Molecular Interaction Platform, Paris, France; Sorbonne Université, US PASS, Plateforme Post-génomique de la Pitié-Salpêtrière (P3S), 75013 Paris, France; Sorbonne Université, CNRS, Inserm, Institut de Biologie Paris-Seine, IBPS, F-75005, Paris, France

**Keywords:** microalgae, ATP synthase, Tara Oceans, chloroplast, environmentally-informed phenotyping

## Abstract

Microalgal metabolism relies on their chloroplasts, and involves both nucleus and plastidial-encoded proteins of various evolutionary origins. The plastidial ATP synthase complex is a key player in photosynthesis, and has been extensively studied in plants. However, our knowledge in other photosynthetic eukaryotes remains limited, despite their importance in marine environments. Here, we report the characterisation of a novel homologue of the F-type ATP synthase alpha subunit, hereby named xATPA, widespread in microalgae but absent from other photosynthetic organisms. Comparisons of xATPA sequences and predicted structures revealed a specific feature, the bump domain, and highlighted the absence of an ATP-binding site. We assessed xATPA prevalence in microalgae in the global ocean using environmental data from *Tara* Oceans, with a particular focus on diatoms, and demonstrate that its expression is associated with polar summer conditions. Using a reverse genetic approach in the model diatom *Phaeodactylum tricornutum*, we show that xATPA_Pt_ has a plastidial localisation, and that xATPA KO mutants exhibit growth deficiencies in a combination of low temperature, low salinity and constant light, consistent with environmental analysis. Surprisingly, both RNAseq and physiological assays suggest that xATPA is not involved in ATP synthase functions. On the other hand, xATPA interacts with other F1 ATP synthase subunits in vitro, which we suggest forms transient unassembled complexes. This study hence represents a comprehensive analysis of a novel protein from the environment to the lab, and reveals a new player in the plastidial physiology of eukaryotic microalgae.

## Introduction

Eukaryotic microalgae are a functional group of phylogenetically diverse unicellular photosynthetic organisms, which can be found in marine, freshwater and terrestrial environments (1). Their chloroplasts have convoluted evolutionary origins and result from secondary, tertiary or higher endosymbioses with other eukaryotic microalgae, serial plastid loss or replacements, transient acquisition, etc (2, 3). Hence, chloroplast functions in microalgae are very diverse and are supported by a mosaic of proteins from various evolutionary origins, for example, acquired horizontally through endosymbiotic gene transfer (EGT) (2, 4). Such events rely on the integration of endosymbiont - derived genes within the nuclear genomes, and the subsequent retargeting of their protein products to the chloroplast (2, 4, 5). Although main plastidial pathways described in plants are generally conserved in other photosynthetic organisms, plastid proteomes of microalgae also contain a high number of actors with unknown functions (4–6).

Such an instance was described in the proteomic analysis of the plastid of the microalga *Euglena gracilis* (Euglenophyta, secondary green chloroplast) (4, 5), in which two homologues of the F-type ATP synthase α subunit were reported: a canonical protein ATPA encoded in the plastid genome, and a divergent protein, that we here name “xATPA”, which is nucleus-encoded. In the chloroplast, the ATP synthase complex is embedded within the thylakoid membrane (F_0_ component). Plastidial ATP synthase exhibits reversible ATP synthesis/hydrolysis activities, but with the stromal F_1_ component mainly catalysing ATP production in the stroma at the end of the light-dependent reactions of photosynthesis (7). Because the ATP synthase complex has been extensively studied in plants, but with no description of xATPA, its discovery in a microalga was surprising and called for further investigations in other eukaryotes.

In this study, we conducted a multi-level approach aiming to decipher the functions of this new xATPA protein. We show that xATPA is found in a diverse range of microalgal organisms, but absent from cyanobacteria, red algae and land plants, with an evolutionary history potentially marked by EGT. Moreover, xATPA sequences and structural predictions reveal that these proteins differ from canonical α subunits by the presence of a specific feature, henceforth called the bump domain, and by the absence of the conserved ATP-binding P-loop motif (7, 8). Using environmental data from *Tara* Oceans (9, 10), we confirm that *xATPA* is expressed in wild microalgae in the global ocean, and is mostly associated in diatoms with polar summer conditions. We explore the biological functions of xATPA in microalgae by conducting a reverse genetic approach in the model diatom *Phaeodactylum tricornutum* (6, 11). We confirm that the xATPA*_Pt_* protein (Phatr3_J31465) is targeted to the chloroplast, and that its functional knock-out causes strong growth deficiencies in a combination of environmentally relevant conditions. Unexpectedly, RNAseq and physiological analyses of KO strains suggest that xATPA*_Pt_* is not involved in ATP synthase functions, whereas experimental assays conducted on purified proteins demonstrate that xATPA*_Pt_* can bind other ATP synthase stromal subunits and may be part of a transient unassembled F_1_ complex in the chloroplast. Together, our work suggest that xATPA is a new actor in the regulation of the plastidial physiology in microalgae, and may participate in their environmental success in marine ecosystems.

## Results

### The pan-microalgal xATPA protein is a divergent homologue of the canonical F-type ATP synthase α subunit

We sought to assess the occurrence of xATPA across the Tree of Life using a custom dataset of more than 700 publicly available genome and transcriptome resources encompassing the broad diversity of eukaryotes (12, 13) (Dataset S1). At first, a set of seed xATPA sequences was identified via a top-down approach, as a distinct ATPA-like clade (xATPA) within phylogenetic trees combining all ATPase subunits found in *E. gracilis* and a selection of eukaryotic transcriptomes. These sequences subsequently served as queries for targeted BLAST searches. In total, 222 xATPA homologues were identified based on sequence similarity, with a mean 70 kDa size and a striking prevalence in photosynthetic species (Fig1A, S1). Moreover, xATPA was never detected in cyanobacteria nor in Archaeplastida (land plants, red algae, glaucophytes. . .) with the exception of Chlorophyta, and is absent from differentiated multicellular algal species (Fig 1A, S2). Hence, xATPA is found in unicellular eukaryotic microalgae, with a presence in Chlorophyta, Euglenophyta, Dinoflagellata, Haptophyta, Chlorarachniophyta, Cryptophyta and Heterokontophyta (pelagophytes, dictyochophytes, pinguiophytes and bolidophytes and diatoms) (1, 3). However, xATPA is found to be absent in several microalgal groups within Heterokontophyta, such as Eustigmatophyta or Chrysophyta, indicating that its functions are not essential in these taxa (Fig 1A, S2). Surprisingly, it is present in several species of labyrinthulomycetes but never detected in other heterotrophic stramenopile protists (oomycetes, *Blastocystis spp.*, etc.) (Fig S2). Hence, xATPA functions may not be strictly related to chloroplast-specific metabolism (Fig 1A, S2). On the other hand, the xATPA phylogeny suggests that its evolutionary history is related to chloroplast transfers within eukaryotes (3). Indeed, tree topology does not follow the species phylogeny, and several EGTs can be identified from the topology, e.g. the green origin of the *E. gracilis* xATPA or the haptophyte origin of its fucoxanthin dinoflagellate homologues (Fig 1A, S2, S3) (3, 5, 14). Moreover, xATPA homologues form a monophyletic group in our tree topology, separated from other mitochondrial, plastidial and bacterial F-type α subunits (ATPA) (Fig 1A, S3). Such distinction between xATPA and ATPA remains conserved when β subunits (ATPB) are included as outgroup, and internal BLAST confirms a significantly higher xATPA identity with ATPA (25.9%) than with ATPB (20.6%, p = 1.9 x 10^−16^, two-tailed paired t-test, df = 221) (Fig S4).

**Fig. 1.**
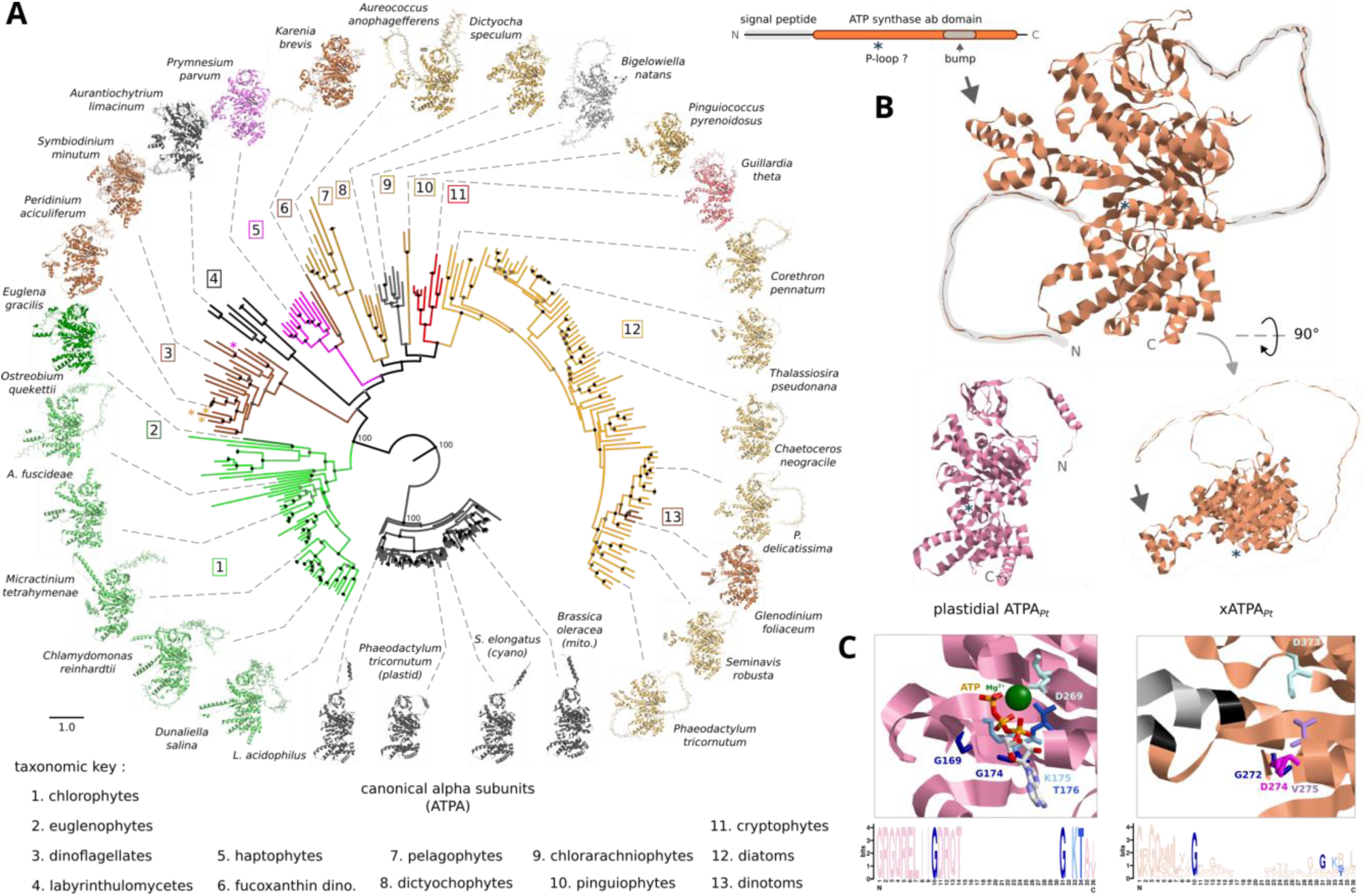
Diversity and characteristic features of xATPA. A: Phylogenetic tree of ATP synthase domains and predicted 3D structures. The tree was built using the 222 identified xATPA sequences, along with canonical ATPA subunits from diverse origins. Branches are coloured based on their taxonomy, the scale bar represents the average amino acid mutation rate, bootstrap branch supports higher than 75 are represented at each node and coloured based on their value: 100 = black, >90 = dark grey, >75 = light grey. In dinoflagellates, stars indicate species that have two xATPA copies, and their colours correspond to the taxa in which the second copy is found. 3D structures are coloured based on their taxonomy, and the name of the corresponding species and its position within the tree are indicated. For xATPA, ATP synthase domains are shown in ribbons while signal peptides are represented in stars, when present. Some targeting sequences were shortened for visualisation purposes. All predicted structures have a predicted template simulating (pTM) above 0.5. B: Predicted structures of the *P. tricornutum* xATPA (orange, UniProt: B7FRE6, average pLDDT = 76%) and plastidial ATPA (pink, UniProt: A0T0F1, average pLDDT = 91%). Schematic xATPA primary structures is represented on top. Position of the P-loop motif is indicated by asterisks on the 3D models. C: Close-up on the ATP binding pocket of the crystal structure of ATPA bound to ATP (pink, PDB: 1E79 chain C) and the predicted structure from *P. tricornutum* xATPA (orange, UniProt: B7FRE6). The bump domain is coloured in grey, flanking amino acids are shown in black. Amino acids are numbered based on their position in each protein sequence. The P-loop motif (GxxxxGKT) and magnesium-stabilising aspartate are shown in blue. xATPA: amino acids positioned at the same place as the ATP-interacting ones in ATPA are shown. Sequence logos show ATPA and xATPA alignments around the P-loop motif.

Sequence alignments and AlphaFold 3 structural predictions clearly show the divergence of xATPA from the canonical ATPA subunits (Fig 1A,B, S3) (15). Although xATPA proteins are predicted to conserve the characteristic structure of F-type α subunits (Fig 1A, 1B) (7), all identified proteins share similar peculiarities. First, they have an unfolded N-terminal sequence which is likely to be a targeting sequence involved in xATPA subcellular localisation (Fig 1A, 1B, S5). Indeed, intracellular targeting predictions indicate an intracellular localisation in either mitochondria or chloroplasts (Fig S5) (16, 17), which possess F-type ATP synthase complexes. Second, xATPA proteins consistently bear at the same position a small domain of about 60 amino acids protruding from the central domain, that we call here the bump domain (Fig 1A, 1B, S6). This domain is positioned between the alanine 444 and the valine 499 in the *P. tricornutum* xATPA protein (Fig S6). However, bump domains show strong differences in length, sequence and structural predictions between green-related and SAR-related xATPA, although sequence conservation is very poor within these two groups (Fig 1A, S6). Third, xATPA homologues do not consistently exhibit the very conserved ATP-binding P-loop motif that is present in all canonical α subunits (8), suggesting that its biological functions do not involve nucleotide binding (Fig 1C). Hence xATPA functions remain elusive, and might not overlap with the canonical functions of mitochondrial or plastidial α subunits.

### Microalgal *xATPA* metatranscript abundances are associated with specific environmental conditions in diatoms

Since xATPA is extensively present in eukaryotic microalgae, we decided to investigate its occurrence in these organisms in the wild. To do so, we searched for xATPA sequences in *Tara* Oceans data (9, 10), which combines both metagenomic and metatranscriptomic information from water samples collected in more than 200 locations across the global ocean, at different sampling depth and for various size fractions, along with corresponding environmental parameters (9). Wild xATPA sequences were then curated and taxonomically classified by phylogenetic reconciliation to the cultured species topology (18) (Fig S7). Overall, *xATPA* metagenes and metatranscripts were detected for chlorophytes, cryptophytes, dinoflagellates, haptophytes, diatoms and other heterokontophyte microalgae, and found to be ubiquitous in all these groups (Fig 2A, S8, S9). However, *xATPA* metatranscript abundances were found to be associated with several environmental parameters, with a strong negative correlation to temperature and salinity (Fig 2B, 2C, S9), suggesting a function of the xATPA proteins in these specific conditions. Due to their prevalence in marine ecosystems (6, 10), we further focused on diatoms to conduct in-depth analyses. In this group, *xATPA* metatranscript abundances are predicted to be higher in polar conditions, which are characterised by combinations of low salinity and temperature and long days in summer. Moreover, both salinity and temperature are found to be the best two predictors for diatom *xATPA* metatranscript abundances (random forests regression model, R^2^ = 0.82) (Fig 2A, S10). In the same way, a principal component analysis confirms the negative correlation between *xATPA* metatranscript abundances in diatoms and temperature and salinity, and its very low correlation with nutrient content (Fig 2B). Together, these results hint at the specific functions of xATPA in such environmental conditions, subsequently referred to as “polar summer” conditions, that we simulated by the combination of low temperature, low salinity and continuous light.

**Fig. 2.**
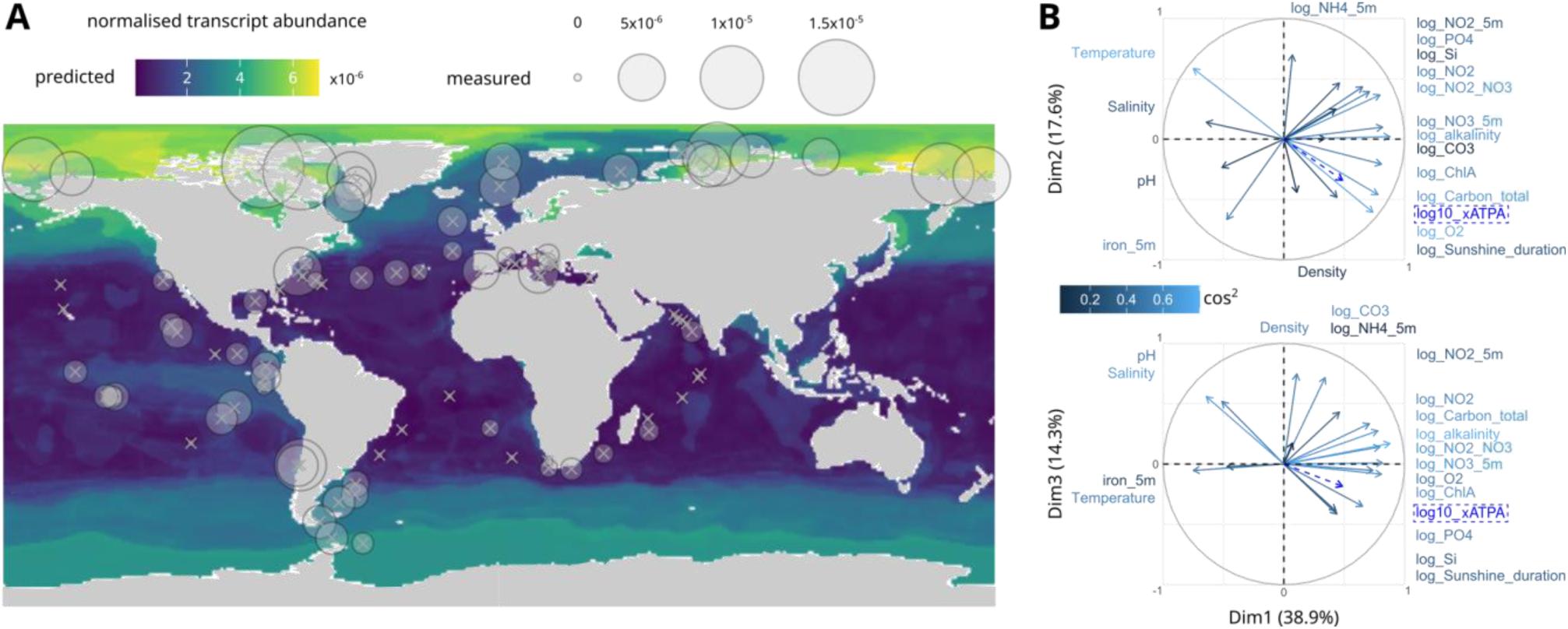
Diatom *xATPA* metatranscript abundances are associated with polar summer conditions in the wild. A: Predicted diatom *xATPA* metatranscript normalised abundances at the surface are represented by a colour gradient, while measured diatom *xATPA* metatranscript normalised abundances at the surface are shown as circles for each station. A random forests regression method was used to predict abundances based on measured values and global oceanographic data (R^2^ = 0.82). Normalised abundances are the log10-transformed mean values calculated within each station in surface samples. B: PCA correlation circles in the first 3 dimensions, with explained variance for each axis is shown as percentages. Computation was performed on *Tara* Oceans environmental variables, with log10 transformation when the normal distribution was not met. Log10-transformed mean diatom *xATPA* metatranscript normalised abundances at the surface in each station were used as supplementary variable (bright blue, dashed lines). Arrow colours and names represent the cos2 value of each environmental parameter.

### Reverse genetic analyses in *P. tricornutum* reveal the essential rôle of the plastidial xATPA protein for growth in environmentally relevant conditions

In order to explore the biological functions of xATPA in microalgae, we conducted a reverse genetic approach in the model diatom *P. tricornutum* (6, 11). We first generated a transgenic strain expressing a chimeric construct made of the endogenous xATPA*_Pt_* protein (Phatr3_J31465, monoexonic gene) with a C-terminal fusion to the eGFP, under the strong Fcp promoter (Fig 3A, S11) (6, 11). Observation of the eGFP fluorescence signal in the cells using confocal microscopy showed a colocalisation with chlorophyll autofluorescence (Fig 3A). This result indicates a plastidial localisation of xATPA in *P. tricornutum*, consistent with previous observations in *E. gracilis* via proteomics of isolated cell fractions (5).

**Fig. 3.**
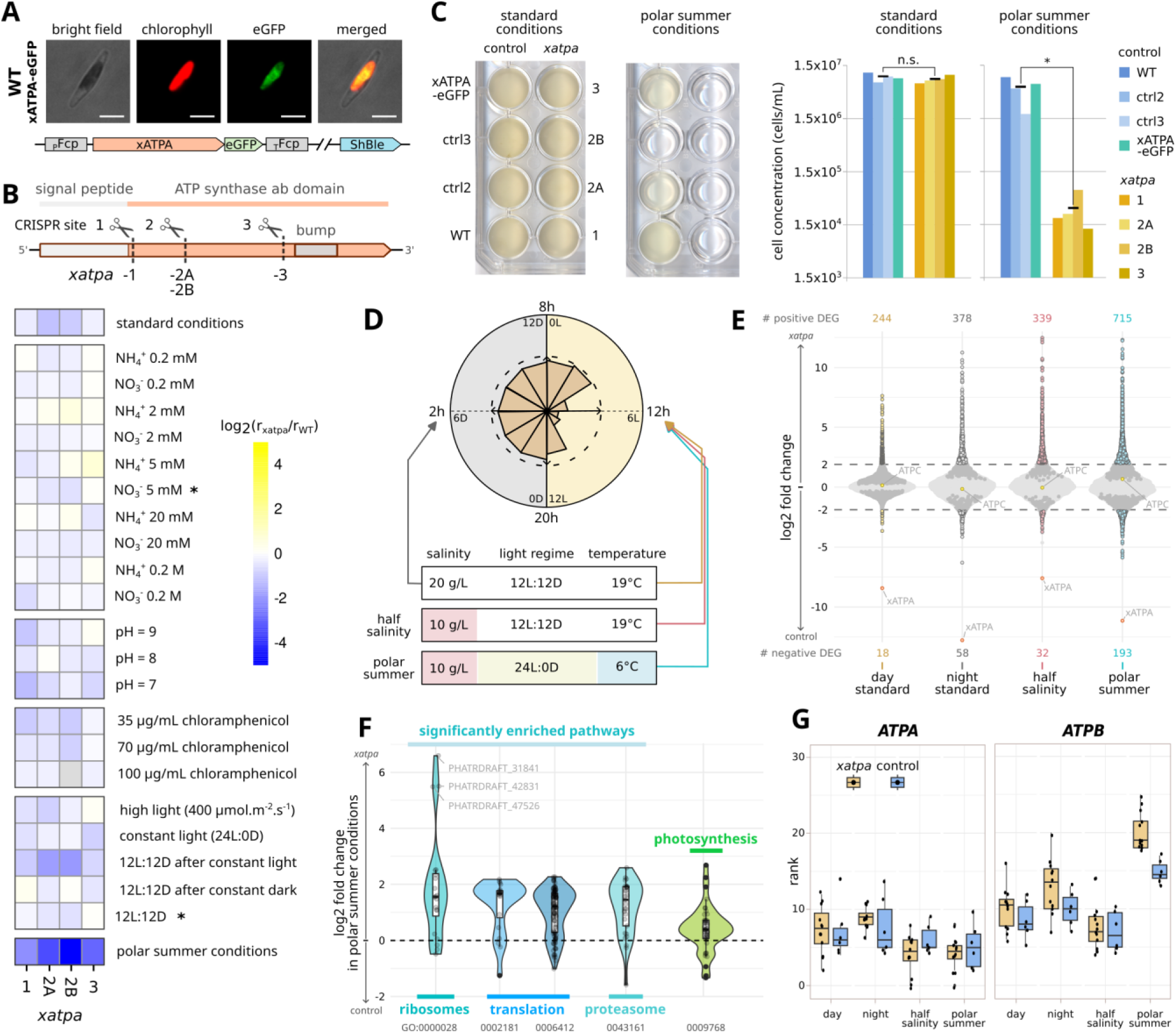
*xATPA* knock-out alters *P. tricornutum* growth and physiology in polar summer conditions. A: Plasitidial localisation of overexpressed xATPA-eGFP fusion protein in wild type *P. tricornutum* . The scale bars represent 5 μm. The genetic construct is presented at the bottom of the image strip. B: Four *xATPA* KO strains (*xatpa*) were generated using the CRISPR-Cas9 system. Control strains *ctrl* were transformed with HA-hCas9 and ShBle resistance plasmids without sgRNA. (Top) Positions of the sgRNA-Cas9 target sites are indicated on the WT *xATPA* gene. All *xatpa* strains are independent and genetically non-identical, and possess frameshift mutations leading to premature stop codons and encoding truncated proteins. (Bottom) Growth curves were performed in various conditions, and relative growth rates of *xatpa* strains compared to the WT were calculated. In the heatmap, columns correspond to *xatpa* strains, lines correspond to the specific growth condition indicated on the right, asterisks correspond to standard conditions. C: Growth assays in 24-well plates, and corresponding final cell concentrations. Two-tailed Wilcoxon tests were performed to assess the significance of mean differences between different control and *xatpa* strains (standard conditions : p = 0.49; polar summer conditions : p = 0.029). D: RNAseq was performed on *ctrl* and *xatpa* cultures in exponential phase, in four different conditions. The sampling plan and culture conditions are represented by the diagram. *xATPA* RNA levels throughout the day as measured by qPCR are represented by the circular plot, where the dashed line indicates the normalised maximum RNA level. The time of the day is indicated on the outside of the circle, light:dark time is indicated on the inside. Each strain is represented by three biological replicates, with two *ctrl* and four *xatpa* strains tested. E: Differential gene expressions were computed in R using the DESeq2 package, with genotype, growth conditions and their interactions as comparison factors (19). Sample conditions are indicated on the x axis, and log2 fold change values between *xatpa* and *ctrl* strains are represented on the y axis for each gene. Genes are considered differentially expressed when their absolute log2 fold change is above 2 and below the 0.05 adjusted p-value threshold. Genes below this threshold but with a lower absolute log2 fold change are represented in dark grey, and other genes are represented in light grey. *xATPA* (Phatr3_J31645) is represented in orange, nucleus-encoded plastidial *ATPC* (Phatr3_J20657) in yellow. The number of differentially underexpressed (bottom) and overexpressed (top) genes in *xatpa* strains are indicated for each condition. F: Distribution of log2 fold change values in the comparison between *xatpa* and *ctrl* strains in polar summer conditions, within a selection of pathways. Significant enrichments were determined by gene set enrichment analyses. Gene Ontology IDs are provided for each pathway. G: Comparison of ranked expression of the plastidial genes *ATPA* and *ATPB* between *xatpa* and *ctrl* strains, for each condition. Plastidial genes were ranked based on their gene counts, with a high rank value corresponding to a high gene count.

With the hypothesis that high *xATPA* metatranscript abundances in the wild reflect a functional importance of the xATPA protein in diatoms, we assessed the impact of various growth conditions on xATPA knock-out mutants of *P. tricornutum*. Using the CRISPR-Cas9 system (6, 20), we obtained four independent *xATPA* KO strains (*xatpa*) by targeting three different CRISPR sites on the *xATPA_Pt_* gene, which lead to both reduced *xATPA_Pt_* RNA levels and shortened proteins (Fig 3B). Despite numerous growth conditions tested, including different nitrogen sources and concentrations, starting medium pH, chloramphenicol (inhibitor of organellar translation) concentrations, several light regimes and intensities, *xatpa* strains showed similar growth phenotype compared to WT strains or control (*ctrl*) transgenic lines transformed with same vectors but without sgRNA (Fig 3B, S12). However, in coherence with environmental data, *xatpa* strains exhibited a dramatic reduction of their growth abilities when cultivated in polar summer conditions, i.e. in half salinity medium at 6°C and under constant light, while remaining viable (Fig 3B, 3C, S13). On the other hand neither control vector nor xATPA*_Pt_*-eGFP overexpression strains showed alteration in their growth phenotype in both standard and polar summer conditions (Fig 3C). Moreover, individual stresses (temperature, salinity and light regime) have lower impact on the growth of *xatpa* strains than all three combined (Fig S14). Hence, the polar summer condition appears as the best setup to study this functional knock-out.

Because the presence of active sgRNA-Cas9 in *xatpa* mutants prevents their complementation with the endogenous *xATPA_Pt_* gene, we generated complemented strains using a labyrinthulomycete *xATPA_laby_* gene (*Aurantiochytrium sp.* RCC893) bearing the xATPA*_Pt_* targeting sequence and fused to the eGFP under the control of the strong NR promoter (Fig S15) (6, 21). However, complemented strains showed similar impairment in their growth abilities in polar summer conditions as *xatpa* strains (Fig S15), suggesting more complex regulations underlying xATPA*_Pt_* biological activities or poor functional conservation between homologues.

It is notable that *xatpa* strains do not present alterations in their Fv/Fm ratio in standard conditions or when transferred to polar summer conditions (Fig S16) (22). Moreover, *xatpa* strains do not exhibit increased sensitivity to light-induced stresses, such as high light intensities or exposure to UV light (Fig S12, S16) (23, 24). Since other subunits KO were reported to alter light-related metabolism in microalgal organisms (25, 26), our results suggest that the xATPA*_Pt_* protein may not participate in the function of the plastidial ATP synthase complex in *P. tricornutum*.

### RNAseq data show that *xATPA_Pt_* KO impacts pathways related to protein metabolism, but not ATP synthase-related pathways

Using RNAseq analyses of *xatpa* and *ctrl* strains in various culture conditions, we sought to unravel the biological functions of xATPA in *P. tricornutum*. Since *xATPA_Pt_* exhibits a circadian expression pattern, with high RNA levels during the dark phase and minimal levels in the middle of the light phase (Fig 3D, S17), we analysed the impact of *xATPA_Pt_* KO on transcript contents at these two time points in standard culture conditions (Fig 3D, S17). We also used the middle of the light phase (day mid-point, 6h after light onset in 12h light/12h dark cycle) as the reference time point for half salinity and polar summer culture conditions (Fig 3D). Strong depletion in *xATPA* RNA levels in *xatpa* strains was confirmed by these data (Fig 3E).

Out of the 12 233 gene models of the v3 *P. tricornutum* genome (11), 18 and 244 genes were found to be down and upregulated (respectively) between *xatpa* and *ctrl* strains in standard conditions at day mid-point, with gene set enrichment analysis (GSEA) showing an enhanced expression of pathways related to proteasome, ribosomes and translation (Fig 3E, S18) (19). In the same way, *xatpa* is associated with the upregulation of 378 genes enriched in proteasome-related and 339 genes enriched in ribosome-related pathways, in standard conditions at night and half salinity at day mid-point, respectively (Fig 3E, S18). Interestingly, when *xatpa* and *ctrl* strains were compared in polar summer conditions, 193 and 715 genes were found to be down and upregulated (1.5 and 6% of total nuclear genes, respectively), and the same enhanced expression of proteasome, ribosomes and translation pathways was detected by GSEA (Fig 3E, 3F, S18). As suggested by our previous results, *xATPA_Pt_* KO did not affect the expression of photosynthesis-related genes, neither in polar summer nor other culture conditions (Fig 3F, S18). Moreover, the only nucleus-encoded plastidial ATP synthase subunit (γ subunit, *ATPC*, Phatr3_J20657) (11, 26, 27) was never found to be differentially expressed between *xatpa* and *ctrl* strains, in any conditions (Fig 3E).

To better understand the physiological consequences of polar summer conditions on *P. tricornutum*, we compared RNAseq data from *ctrl* strains in standard and polar summer conditions at day mid-point. Interestingly, this analysis resulted in 247 downregulated and 620 upregulated genes (2 and 6% of total nuclear genes, respectively), and GSEA indicated an enhanced expression of translation and ribosome pathways, and reduced expression of light-harvesting pathways (Fig S18). Such results correspond to what was consistently observed between *ctrl* and *xatpa* strains, suggesting that *xATPA_Pt_* KO may induce a physiological response comparable to the one triggered by polar summer conditions (Fig 3F, S18).

Moreover, weighted gene correlation network analysis (WGCNA) conducted on our data clearly demonstrated an association between *ATPC* and genes involved in energetic metabolism, as expected for an F-type ATP synthase subunit, while *xATPA_Pt_* did not (Fig S19) (28). Once again, such results strongly suggest that xATPA*_Pt_* is not involved in the function of the plastidial ATP synthase complex. Ranked analysis of read counts of plastid-encoded genes shows higher RNA levels of the β subunit (*ATPB*) and of its 5’ UTR/upstream *ycf3* gene in polar summer conditions compared to other samples, this phenotype being more intense in *xatpa* strains (Fig 3G, S20) (27). On the contrary, the plastidial α subunit gene (*ATPA*) shows low and stable ranks across conditions (Fig 3G). This suggests a possible link between xATPA*_Pt_* functions and *ATPB* expression in the chloroplast (25), associated with polar summer conditions.

### xATPA*_Pt_* interacts with the β subunit and may form a transient incomplete F_1_ complex in the chloroplast

Even though xATPA*_Pt_* may be not involved in ATP synthase functions in photosynthesis, its protein sequence and predicted 3D structure suggest that it may have similar molecular properties as the canonical α subunit. In order to assess for their functional redundancy, we used a *Saccharomyces cerevisiae* BY4147 yeast strain knocked-out for its mitochondrial F-type α subunit (Δ*ATP1*), and compared the impact of the heterologous expression of xATPA*_Pt_* and plastidial ATPA*_Pt_* bearing the ATP1 mitochondrial targeting signal. Surprisingly, differences in growth abilities of transformed strains suggest that xATPA*_Pt_* and canonical α subunits do not share similar properties *in vivo* (Fig S21). However, *in silico* interaction between xATPA*_Pt_* and other plastidial F_1_ subunits predicts that it is able to bind to a β subunit and to form heterodimers with other F_1_ components (Fig 4A, S22). In these complexes, the bump domain is predicted to prevent the binding of the last β subunit, which would limit the assembly to a 2α:2β:1xATPA heteropentamer inserted on the γ subunit (ATPC, Fig 4A). Such interactions are also predicted to occur with xATPA proteins from other distantly related microalgae, suggesting a functional conservation of xATPA properties across eukaryotes (Fig S22).

**Fig. 4.**
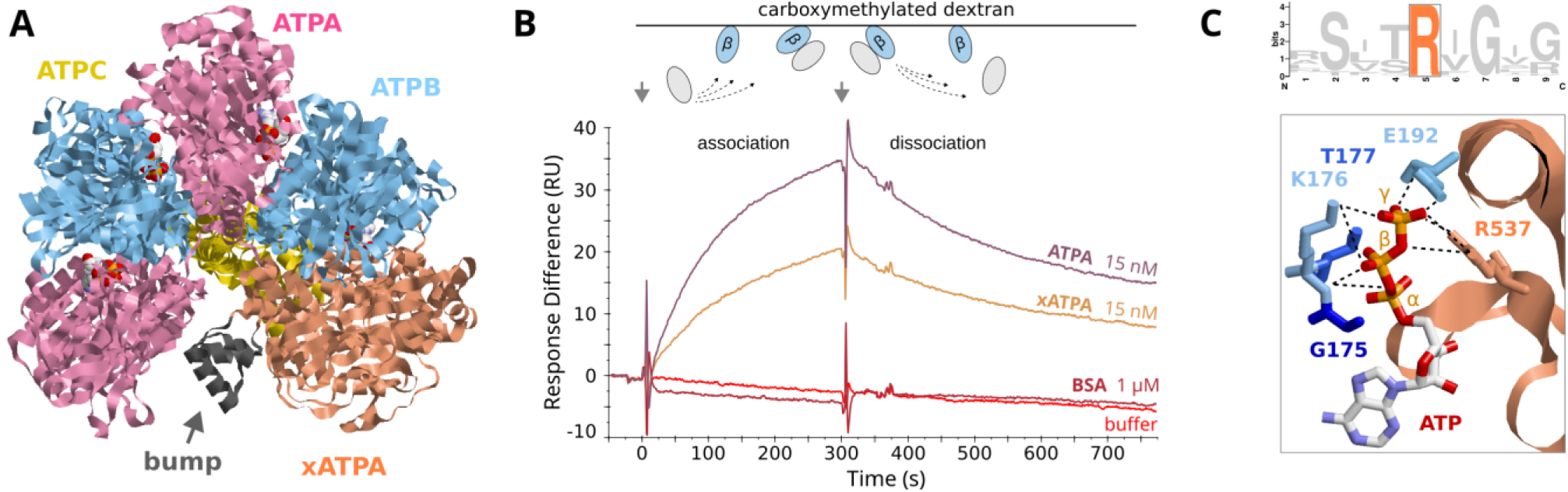
The xATPA protein interacts with other F_1_ ATP synthase subunits. A: AlphaFold 3 predicted structures and interactions of *P. tricornutum* plastidial ATPB (blue, UniProt: A0T0D2), plastidial ATPA (pink, UniProt: A0T0F1), plastidial ATPC (yellow, UniProt: Q84XB4) and xATPA (orange, UniProt: B7FRE6), pTM = 0.80, ipTM = 0.77. The xATPA bump domain is shown is grey, and prevents the binding of a third ATPB subunit. B: Surface Plasmon Resonance assay profiles using immobilised CBP-ATPB-6His. A diagram of the process is shown on top. First arrow (t=0s, association phase): buffer flow with a fixed analyte concentration, second arrow (t=300s, dissociation phase): buffer flow without analyte. Colours correspond to different conditions, red: buffer only (baseline), dark purple: 1 μM BSA, orange: 15 nM TwinStrep-xATPA-6His, pink: 15 nM TwinStrep-ATPA-6His. For a same concentration, TwinStrep-ATPA-6His shows stronger association with the β subunit compared to TwinStrep-xATPA-6His. C: (Top) Sequence logo of xATPA ATP-binding arginine region, with the conserved arginine highlighted. (Bottom) Close-up on the ATPB ATP-binding pocket in a predicted *P. tricornutum* ATPB:xATPA dimer (pTM = 0.83, ipTM = 0.86). For ATPB, only important amino acids are represented. Phosphate-interacting amino acids are shown as sticks, blue: ATPB, orange: xATPA. Amino acids are numbered based on their position in each sequence, and phosphate group order is shown. Black dashed lines represent weak interactions between ATP and amino acids as predicted by AlphaFold 3. The Mg+2 ion is not shown.

To experimentally confirm whether xATPA*_Pt_* can bind its putative β subunit partner, we expressed and purified double-tagged *P. tricornutum* proteins in *E. coli*, to obtain the full length TwinStrep-xATPA-6His as well as the plastidial TwinStrep-ATPA-6His and CBP-ATPB-6His subunits. In vitro interaction followed by pull-downs on either partners demonstrated their ability to bind together (Fig S23). We also sought to identify other xATPA protein partners in *P. tricornutum* by performing a pull-down on purified xATPA*_Pt_* with whole cell extracts, followed by mass spectrometry (Fig S23). Interestingly, both ATPB and ATPC proteins were identified as xATPA *_Pt_* protein partners (Fig S23), supporting its integration within an incomplete F_1_ complex. Finally, Surface Plasmon Resonance measurements revealed that xATPA*_Pt_* has lower affinity for ATPBP t than ATPA*_Pt_*, with an *in vitro* dissociation constant of the xATPA:ATPB dimer five times higher than for ATPA:ATPB (1.2x10^−8^±0.1 and 2.3x10^−9^±0.49, respectively) (Fig 4B, S24) (29). Thus, xATPA is a weak ATPA competitor for ATPB binding.

Within the F_1_ complex, ATP synthesis or hydrolysis require the cooperation of both α and β subunits at the ATPB ATP-binding site (7, 30). This process involves the guanidino group of a conserved ATPA arginine (7, 30), which is also well conserved in all xATPA proteins (Fig 4C). This observation suggests that xATPA may act in the catalytic regulation of plastidial β subunits, by either inhibiting or enhancing its ATPase and/or ATP synthase activities.

## Discussion

### Presence of xATPA in eukaryotic microalgae

In this study, we sought to understand the functions of xATPA, a distant homologue of the F-type ATP synthase α subunit. Our work reveals that xATPA is widespread across the tree of eukaryotes, with a predominant association with microalgal organisms, and subcellular targeting in organelles containing F-type ATP synthase complexes. Due to their sequence and predicted structural homologies with canonical α subunits, we hypothesise that *xATPA* originates from the duplication and subsequent insertion within the nuclear genome of a nucleus-encoded mitochondrial or plastid-encoded *ATPA* gene, followed by the insertion of the bump domain coding sequence. *xATPA* genes may have then been predominantly acquired through EGT during chloroplast biogenesis in eukaryotic microalgae (2, 4, 5, 14). Although they share the same position within all xATPA sequences, differences in the bump domains between green and SAR-related xATPA suggest either an independent acquisition and convergence in these two groups (non-homologous), or a common origin followed by strong divergence (homologous). However, the presence of several genes originating from a putative ancestral green endosymbiont have been reported in diatoms and in other stramenopiles (4, 31, 32), which may favour the scenario of an first origin of *xATPA* in Chlorophyta, followed by its horizontal transfer and divergence in SAR microalgae.

In addition to their identification in microalgal genomes and transcriptomes, *xATPA* genes and transcripts are also detected in environmental samples. Moreover, the xATPA protein was detected in several proteomic studies conducted in various microalgae, such as the diatoms *P. tricornutum* (33) and *Thalassiosira pseudonana* (B8LDX7 & B8LDX8, split gene annotations) (34), or the green algae *Dunaliella salina* (DsB g19279) and *D. tertiolecta* (Dte g04731) (35), as well as in *Chlamydomonas reinhardtii* (Cre10.g419050) (34, 36). The diel expression pattern of xATPA is also conserved across microalgal groups, as reported in the diatom *Seminavis robusta* (Sro536_g162070)(37) and the green algal model *C. reinhardtii* (38). In the latter, xATPA is known as ATP1B, and is annotated as an isoform of the mitochondrial canonical α subunit ATP1A (39, 40). However, to our knowledge, ATP1B has neither been experimentally localised in mitochondria nor shown to be involved in their metabolism, while its diel expression is synchronised with canonical plastidial ATP synthase subunits (38).

### Molecular mechanisms underlying xATPA functions in microalgae

Our results demonstrate that the xATPA protein from *P. tricornutum* interacts with the β subunit and may integrate plastidial incomplete F_1_ complexes. Interestingly, a similar non-catalytic, 70 kDa, F_1_-associated protein cross-reacting with anti-beta/alpha subunits antibodies was identified from *C. reinhardtii* thylakoid fractions about 35 years ago (41). Such similarities lead us to consider that this protein may be in fact the *C. reinhardtii* xATPA homologue (ATP1B), but with a plastidial localisation. It can be hypothesised that xATPA acts as regulator of the catalytic activity of the plastidial β subunits or of the incomplete F_1_, which could be tested by the isolation and enzymatic characterisation of these complexes (41).

Despite being a common feature of all xATPA proteins, the roles of the bump domain remain to be explored. In addition to its role in steric blocking within the F_1_ complex, this domain could also act as a plateform to recruit other protein partners, or participate in the regulation of the assembly or of the catalytic activity of the complex (7, 25). Further investigations would require both *in vivo* and *in vitro* approaches in order to determine whether this domain is necessary or sufficient for xATPA biological activities.

In chloroplasts, the expression of ATP synthase subunits is tightly regulated (7, 25, 42), and functional KO of the nucleus-encoded plastidial ATPC subunit in both *C. reinhardtii* (36) and Cyclotella cryptica (diatom) (26) have been reported to induce lower expression levels of its plastidial F_1_ partners, but with no effects on their xATPA homologues (26, 36). Moreover, *C. reinhardtii* xATPA protein levels are not affected by mutations reducing plastidial levels of α and β subunits (41, 43). Hence, xATPA expression in microalgae seems not to be dependent on the expression of other plastidial F-type ATP synthase components, in contrast to canonical subunits. Moreover, our results strongly suggest that xATPA functions are not related to photosynthetic metabolism in *P. tricornutum* .

In *C. reinhardtii*, the expression and assembly of the F_1_ subunits are modulated by a process called Control by Epistasy of Synthesis (CES) (25), in which unassembled α:β subunits regulate their own expression (42), thus ensuring their stoichiometry within the ATP synthase complex. To date, such a mechanism has not been explored in diatoms, but plastidial ATP synthase biogenesis can be hypothesised to rely on similar mechanisms. Based on our results, we propose that xATPA*_Pt_* could be involved in this mechanism, and may regulate the β subunit biogenesis (25). xATPA*_Pt_* could act through its interaction with either accumulating monomeric ATPB or incomplete F_1_ complexes, by altering their transcription, translation or degradation rates. The precise monitoring of these processes in *xATPA* KO strains in *P. tricornutum* would allow to further explore its role in plastid biology.

### Physiological roles of xATPA in *P. tricornutum* exposed to polar summer conditions

In the diatom *P. tricornutum*, *xATPA* KO induces growth impairment in a combination of specific conditions (i.e. low temperature, low salinity and constant light), which reflects its expression pattern observed in diatoms and other microalgal groups in the wild. Our RNAseq data suggest that *xATPA* KO may induce a consistent change in protein metabolism compared to *ctrl* strains, as it is observed when the latter are placed in polar summer conditions. However, the biological mechanisms linking xATPA*_Pt_* molecular properties and the physiological consequences of its functional knock-out on gene expression and growth abilities in polar summer conditions remain elusive. Since protein synthesis and degradation are highly ATP-consuming processes such phenotypes may accommodate fluctuations in ATP levels (44–46), which could be induced for both *xATPA* KO and in polar summer conditions by undefined mechanisms. Thus, comparison of protein translation rates and degradation of these strains by puromycin labelling may provide more insights into the physiological response of *P. tricornutum* to these conditions (47).

In an ecological perspective, xATPA may provide increased acclimation abilities to microalgae in response to changes in their environment. Indeed, its functions may involve translational and post-translational control of cell metabolism, which enable quick and efficient regulation of protein activities (45). Thus, xATPA may provide great evolutionary advantages to microalgae in their natural habitats. This could explain the ubiquity of xATPA across the tree of eukaryotes, as well as its environmental prevalence.

## Acknowledgements

We thank S. Liu for initial plasmid preparation; N. Ruiz Gutierrez for assistance in protein expression; L. Graf for support in bioinformatic analyses. MPR, ANV, VP and RGD acknowledge funding from an ERC Starting Grant (ChloroMosaic, grant number 101039760, 2023-2027) awarded to RGD. ANV and RGD further acknowledge an ANR JCJC (PanArctica, grant number 21-CE02-0014, 2021-2022) awarded to RGD. MPR acknowledges a CDSN doctoral fellowship from the ENS, awarded 2022-2025.

## Materials & Methods

The work presented in this article is adapted from the PhD manuscript “Functional characterisation of the new plastidial ATP synthase subunit xATPA in microalgae : from the molecule to the ecosystem” defended by Mathias Penot-Raquin on 20th March 2026 (1), and available as a [public HAL archive](https://theses.hal.science/tel-05645837). Datasets, scripts, predicted structures, plasmid maps, sequence alignments, raw phylogenetic trees are accessible on this public repository (osf.io/89vm3) and upon request.

### xATPA protein homologues - Structure, Predicted Interactions & Phylogeny

#### Sequences mining in publicly available genomes and transcriptomes

A first set of 27 xATPA protein sequences was initially built starting from the *E. gracilis* sequence. HMM searches were performed on separate alignments of ATPA and ATPB, against a custom database composed of MMETSP transcriptomes (2), in order to retrieve all ATPA/B-like proteins. Manual phylogenetic sorting of the initial hits allowed to identify homologues of the *E. gracilis* xATPA protein, which formed a group distinct from canonical ATPA proteins in the topologies (Dataset S1).

Then, in an early step, xATPA homologues were mined by reciprocal best-hit BLASTp from a custom dataset of 726 genomes and transcriptomes representing the whole diversity of Eukaryotes, as well as Bacteria and Archaea, generated in a previous study (3) and complemted by 107 decontaminated genomes algal genomes from the algallCODE phase II project (4) (Dataset S1). The reference set of 27 xATPA protein sequences described above was used as a first database, and searched by each sequence from our custom genomes and transcriptomes dataset using an e-value threshold of 10^−5^ (3). Best-hit sequences were then used as queries and searched with the same 10^−5^ e-value threshold into the *Phaeodactylum tricornutum* genome version 3 annotation (5). Only the queries able to retrieve the *P. tricornutum* xATPA sequence as a best-hit were conserved for further analyses. These putative homologues were then aligned to the reference xATPA sequences, complemented by ATPA and ATPB sequences from diverse plastidial, mitochondrial and bacterial origins, using MAFFT (- -auto option), and trimmed using trimal (-gt 0.5 option) (6, 7). Neighbour Joining trees were built using built using Geneious v. R.10.09 to iteratively identify homologues, and divergent sequences were removed based on visual curation (i.e. long branches, short or partial sequences). In a later step, most recent sequences published between 2021 and 2025 were individually inspected by BLASTp on NCBI (accessed on 02/09/2025) using xATPA queries specific to the searched taxon.

Alignments were manually inspected to confirm the presence of a bump domain in xATPA homologues, characterised by a gap in canonical ATPA sequences, and to remove very long/short or duplicated sequences using the Seaview v5.0.4 (8) and Taxus v0.3.1 softwares (Taxus). For each taxa, several AlphaFold 3 predicted structures were used to visually validate xATPA identifications, based on the presence of a bump domain and an ATPA-like structure (9). This process resulted in a final set of 222 validated xATPA sequences (Dataset S1).

#### xATPA protein phylogeny

Alignments were performed on Seaview v.5.0.4 using Clustal Omega (8, 10), with a set of mitochondrial and plastidial ATPA sequences as outgroups (Dataset S1). ATP synthase and bump domains were extracted from this alignment using the R package Biostrings v2.74.0 (from aligned amino acid positions 588 to 2171 and 929 to 1133, respectively). Before phylogenetic tree construction, ATP synthase domains were realigned while bump domains were manually curated based on the initial alignment.

Phylogenetic trees were built using IQ-TREE v3.0.1 (11) with -m MFP (best substitution model) and -B 1000 (ultrafast bootstrap replicates) options (12, 13).

#### Organelle targeting predictions

Protein intracellular localisations were predicted using various programmes : ASAFind v1.1.7 (14), HECTAR v1.3 (15), DeepLoc 2.1 (16), MitoFates 1.2 (17), PredAlgo 1.0 (18), SignalP v5.0 (19) and TargetP (20). DeepLoc localisations were found to be representative of the consensus prediction, and only these results were retained for further analyses.

#### Structural analysis of xATPA proteins

X-Ray diffraction structure of the bovine heart mitochondrial ATP synthase F_1_ complex (PDB: 1E79) was used as the reference 3D structure (21). When available, xATPA AlphaFold predicted protein structures were downloaded from the corresponding UniProt entry (Table 1). Remaining structures were obtained using the AlphaFold 3 web server (9). The model_0 3D structures were visualised with Rastop v2.2 (https://www.geneinfinity.org/rastop/).

Sequence logos were generated by the WebLogo server v2.8.2 (22) using the P-loop and ATP-binding arginine domains extracted from the alignments (from amino acids 426 to 462 in the ATP synthase domain alignment, and 1804 to 1814 in the full length alignment, respectively).

### Environmental analyses in *Tara* Oceans

#### xATPA sequence identification in Tara Oceans datasets

Putative xATPA homologues were mined from the eukaryotic MATOU-v1.5 dataset (*Tara* Oceans and *Tara* Polar Circle) (23–25). A HMMer v3.1b2, (https://hmmer.org) search was performed using the ATP synthase Pfam (ATP-synt_ab/PF00006) to identify putative *xATPA* unigenes (26). Their sequences and abundances in every sample were retrieved in both metatranscriptomic and metagenomic datasets (Dataset S2).

Environmental sequences were processed through the xATPA mining and phylogenetic tree building steps described above. Briefly, a BLASTp search was performed using each environmental sequence as a query, in a database made of the initial set of 27 xATPA reference sequences, using an e-value threshold of 10^−5^. Resulting putative xATPA sequences were then searched against the *P. tricornutum* genome version 3 (5), and only sequences able to retrieve the *P. tricornutum* xATPA homologue as a best-hit with an e-value threshold of 10^−5^ were conserved for further analyses. These sequences were then aligned to the reference set of xATPA, ATPA and ATPB sequences and iteratively trimmed based on their position and branch length in Neighbour Joining trees.

The validated environmental xATPA sequences were aligned to the ATP synthase domain HMM profile using the profile alignment option of Clustal Omega available in Seaviewv.5.0.4 (8, 10). Conserved ATP synthase domains were extracted from the alignment using the R package Biostrings v2.74.0. A phylogenetic tree was built with IQ-TREE v3.0.1 (11) using the substitution model Q.PFAM+F+R10 (same as for the ATP synthase domain tree) and -B 1000 (ultrafast bootstrap replicates) option (13), with a topology constrained by the ATP synthase domain tree. Environmental xATPA sequences were taxonomically assigned based on their position in this tree (Dataset S2).

#### Normalisation of environmental xATPA abundances in the different taxa

Metatranscriptomic or metagenomic abundances of environmental xATPA were summed for each sample (combination of station, size fraction and depth) and for each taxonomic group, and normalised by the total metagenomic or metatranscriptomic unigene abundances that are assigned to the corresponding taxon, in the corresponding sample (Dataset S2) (23, 26) :

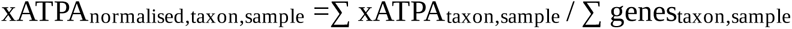

#### Environmental influence on xATPA abundances

*Tara* Oceans environmental and geographical data available for each sample were used to study *xATPA* distribution in the oceans, and to study the Spearman’s rank correlations between *xATPA* abundances and different environmental variables (Dataset S2) (23, 24, 26). Analyses were conducted in R using the packages tidyr, dplyr, ggplot2, maps, ellipse, RcolorBrewer.

For PCA, *xATPA* metatranscript normalised abundances were used as log10-transformed mean values calculated within each station in surface samples. Log10 transformation was performed on variables with non-normal distributions and the missing values of selected envrionmental parameters were imputed by regularised iterative PCA using the missMDA function of the FactoMineR package (27). PCA was performed using *xATPA* abundances as supplementary variable. The packages factoextra, VIM, patchwork and scales were also used.

Predictions of *xATPA* distribution were performed using a random forests regression approach with the R package tidymodels, 80/20 training/testing data ratio, and hyperparameters set to trees=150 and min_n=10. Environmental parameters were downloaded from the World Ocean Atlas 2018, with 1° resolution, and with objectively analysed mean over the 2005-2017 period when available (28). These global environmental parameters were used to replace environmental parameters from the *Tara* Oceans datasets. *xATPA* metatranscript abundances used to train the model correspond to the log10-transformed mean values calculated within each station in surface samples. WOA data and *Tara* Oceans *xATPA* metatranscript abundances were linked based on the sampling stations geographical coordinates. This model was used to predict *xATPA* global abundances from the WOA oceanographic data. Packages vip and pdp were also used.

### Functional study in Phaeodactylum tricornutum

#### P. tricornutum culture conditions

Unless otherwise stated, *P. tricornutum* strains were cultured in 20 mL ESAW supplemented with F/2 nutrients in 25 cm³ flasks wih vented caps. Usual culture conditions (standard conditions) are a temperature of 19°C, an artificial light set at 50 μmol photons.m^−2^.s^−1^ intensity with a 12h light:12h dark cycles, with weekly manual cell resuspension. Solid medium was prepared by mixing equal volumes of ESAW supplemented with nutrients and water, with a final 1% agar concentration, and 100, 50 and 30 μg/mL chloramphenicol, kanamycin and ampicillin, respectively.

Half salinity medium was obtained by a 1:1 dilution of ESAW with pure water, followed by nutrient addition. The polar summer conditions correspond to half salinity medium at 6°C with constant artificial light set at 50 μmol photons.m^−2^.s^−1^ intensity.

Zeocin resistant strains were grown in presence of 100 μg/mL zeocin (InvivoGen) in ESAW. Blasticidin resistant strains (complementation) were grown in presence of 5 μg/mL blasticidin-S (Sigma-Aldrich) in half salinity medium.

#### Plasmid construction and genotyping

Genotyping of the different *P. tricornutum* strains was routinely performed by using 1 μL of culture as DNA template in a 20 μL PCR mix. Oligonucleotides used for gene cloning and CRISPR sites genotyping are described in Dataset S3.

The plasmids necessary to perform the knock-out of xATPA in *P. tricornutum* consist of three types of plasmids (Dataset S3). First, the pU6-sgRNA plasmids allow the expression of single-guide RNAs under the control of the pU6 promoter, with individual plasmids targeting a specific CRISPR sites (Table 2) when associated to the Cas9 endonuclease *in vivo*. Second, the pDEST-HA-hCas plasmid drives the expression of the HA-hCas9 (human codon optimised Cas9 bearing the hemagglutinin epitope in its N-terminus) under the control of the Fcp promoter and Fcp terminator. Third, the pPhaT plasmid confers resistance to zeocin/bleomycin antibiotics via the expression of the ShBle gene under the control of the Fcp promoter and Fcp terminator.

The pPhaT-xATPA-eGFP plasmid was obtained via Gibson assembly in a pPhaT plasmid backbone (Dataset S3). It allows the expression of the fusion protein xATPA-eGFP C-terminal fusion construct under the control of the Fcp promoter and Fcp terminator, as well as zeocin/bleomycin resistance.

Complementation plasmids were obtained via Gibson assembly in a modified pPhaT plasmid backbone, bearing a blasticidin deaminase resistance gene under the control of the Fcp promoter and terminator (29) (Dataset S3). The *xATPA* gene from *Aurantiochytrium sp.* RCC893 was directly amplified from genomic DNA, due to its monoexonic structure in this organism. The xATPA coding sequence was fused with the eGFP at its C-terminus, placed under the control of the NO^−3^ inducible NR promoter and terminator (Phatr3_J54983) (30). The xATPARCC893 target peptide was replaced by the one from its xATPA*_Pt_* homologue to ensure similar subcellular localisation in *P. tricornutum*.

#### Transformation of P. tricornutum

“WT” corresponds to the Pt1.86 - CCAP 1055/1 - CCMP632 *P. tricornutum* strain, and all mutants used in this study were derived from this strain. All *P. tricornutum* transformations were generated by biolistic transformation using the PDS-1000/He system (BioRad), as described in (31). Briefly, the day before bombardment, *P. tricornutum* plates were prepared by plating 5x10^7^ cells from cultures in late exponential exponential phase on solid medium, and microcarriers and 1550 psi rupture disks were sterilised with absolute ethanol. On the bombardment day, 2-5 μg plasmids were coated on tungsten beads using spermidine, and delivered to the cells. Bombarded plates were placed two days in low light to allow cell recovery. After two days cells were harvested, transferred on selective medium, and grown in usual conditions until the apparition of colonies, usually after 4 weeks.

*xATPA* KO strains (*xatpa*) were obtained by co-transformation of three plasmids : pPhaT bearing the BleSh bleomycin/zeocin resistance gene, pU6-sgRNA expressing the single-guide RNA, and pDEST-HA-hCas9 expressing the Cas9 endonuclease. Cas9 target sites were selected using the CRISP-Ex online tool (32). Control strains were obtained by co-transformation of both pPhaT and pDEST-HA-hCas9, but without pU6-sgRNA.

Since individual colonies were mixtures of non-mutated, heterozygous and homozygous cells at the target site, the obtention of homozygous xATPA strains required several iterations of single - colony isolations. To do so, individual colonies were streaked on fresh selective ESAW plates. Each time, gentoyping of single-colonies was performed at the CRISPR target sites by PCR amplification followed by Sanger sequencing (Dataset S3), and subsequent isolations were done until the obtention of homozygous mutants bearing frame-shifting mutations. Colonies with the highest proportion of mutated sequences were identified from Sanger sequencing chromatograms using the Inference of CRISPR Edits (ICE) methodology (33), and were preferentially chosen for the subsequent isolation of xATPA strains.

After transformation of plasmids encoding xATPA fused to the eGFP, single colonies grown on selective media were genotyped with the corresponding Gibson cloning primers to assess for the presence of eGFP.

#### Microscopy

Fluorescence images were obtained at the IBENS Imaging Facility using a LEiCA SP8 confocal microscope, with the following parameters : eGFP excitation = 485 nm, eGFP detection = 500-540 nm, chlorophyll autofluorescence detection = 600-700 nm. WT cells were used as a negative control to confirm detection specificity. Images were false-colored and merged with ImageJ.

#### Growth tests

Growth curves were performed in 25 cm³ flasks wih vented caps, with a starting concentration of 1x10^4^ cells/mL in a final volume of 15 mL medium, without any antibiotics. Cell concentrations were measured either by visual count using Malassez cell or by semi-automatic count using the Luna FX7 cell counter (Logos Biosystems). Growth parameters were then calculated in R by the following procedure : a generalised additive model was first fitted to the experimental data to reduce experimental noise, and smoothed values were then fitted to a Gompertz growth model to estimate the best growth parameters. This procedure was performed in R version 4.4.1 using the packages grofit (34), gam, dplyr, tidyr and ggplot2, pheatmap.

24-well plate growth tests were performed in 2 mL liquid medium with a starting concentration of 1x10^4^ cells/mL, without any antibiotics, and parafilm was used to seal the lid and prevent evaporation. Cultures were grown without agitation, and final cell concentrations were measured using the Luna FX7 cell counter (Logos Biosystems).

UV resistance assays of *P. tricornutum* strains (35) were perfomed by spotting cells in late exponential phase on solid medium, and exposing them to UV light in a Stratalinker UV crosslinker 1800 (Stratagene). Plates were then left for a week in standard culture conditions.

#### Fv/Fm measurements

Fv/Fm measurements were performed on cultures grown in flasks using the QY mode of the AquaPen-P AP 110-P (Photon Systems Instruments), in the middle of the light phase. Cultures were dark-acclimated for at least 5 min, and measurements were performed in the dark at room temperature (approximately 20°C). For cultures grown in cold conditions, dark-acclimation was done in a closed polystyrene box equilibrated at 4°C with ice packs. To avoid signal saturation of chlorophyll fluorescence in dense cultures, 1 mL of culture was diluted in 9 mL PBS at the relevent temperature in a 25 cm³ flask, and dark-acclimated prior to measurement.

For the polar summer acclimation experiment, 20 mL liquid cultures in exponential phase were split in two flasks, and placed either in standard conditions or at low temperature and constant light. Fv/Fm was then measured each day in the middle of the light phase (relatively to the 12L:12D cycle).

#### RNA extraction and RT-qPCR

Cultures in exponential phase (1∼5x10^6^ cells/mL) were harvested by centrifugation and washed with 1 mL PBS, and cell pellets were frozen in liquid N2. RNA was extracted by adding 1 mL TRIzol (Invitrogen) and 200 μL chloroform to the frozen pellet, the mix was vortexed vigorously and centrifuged 2 min at 4°C and 10 000 g. The supernatant containing RNA was carefully taken up and transferred in a fresh RNAse-free tube, 500 μL chloroform was added, the mix was vortexed and centrifuged again. The supernatant was transferred in a fresh RNAse-free tube, mixed with 500 μL isopropanol and left at -20°C for several hours to allow RNA precipitation. After 30 min centrifugation at 4°C and 10 000 g, RNA pellet was washed with 1 mL ice-cold absolute ethanol and centrifuged again 5 min in the same conditions. After removing ethanol, pellet was dried by evaporation, resuspended in 50 μL RNAse-free water. RNA integrity was assessed by electrophoresis on 1% agarose gel, and concentration was quantified using by spectrophotometry with a NanoDrop (Thermo Scientific). Samples were conserved at -20°C.

DNA digestion of RNA samples was performed using RQ1 DNAse (Promega), following manufacturer’s instructions using 2 μg RNA per reaction. Isopropanol-ethanol purification was performed using the same protocol as decribed above. RNA pellets were resuspended in 20 μL RNAse-free water, RNA integrity was assessed by gel electrophoresis, and samples were conserved at -20°C.

For RT-qPCR, retrotranscription was performed using the Maxima First Strand cDNA synthesis kit (Thermo Fisher) or the SuperScript IV kit with random hexamers (Invitrogen) using manufacturer’s instructions. qPCR was performed using the Takyon No ROX SYBR 2X MasterMix blue dTTP kit following the manufacturer’s instructions. xATPA was amplified using the q1, q2 and q3 primer pairs, while the RPS and TBP housekeeping genes were amplified with the corresponding primers (Dataset S3) (36). PCR amplifications were performed with the following progam : 3 min at 95°C, followed by 40 cycles of 10 s at 95°C and 45 s at 60°C (CFX Opus 96 Real-Time PCR system). Technical triplicates were done for each primer used, and xATPA relative expression was calculated using the 2−ΔΔCt method.

#### Biomass production and RNAextraction for RNAseq

Cultures were performed in 75 cm³ flasks wih vented caps, with a starting concentration of 5x10^4^ cells/mL in a final volume of 50 mL medium. For each strain, a unique seed culture in exponential phase was used to create three biological replicate cultures. Cultures were grown for four days in standard conditions. In these conditions, cultures were in exponential phase when harvested (∼1x10^6^ cells/mL). For polar summer conditions, cultures grown four days in half salinity medium were transferred in the middle of the light phase at 6°C, constant light, under a 50 μmol.m^−2^.s^−1^ photon flux, and left three days for acclimation.

Cells were harvested at room temperature by centifugation in 50 mL tubes, washed with 1 mL PBS and centrifuged again in 2 mL tubes. Cell pellets were then flash frozen in liquid N2 and conserved at -80°C. Cultures were always harvested at the same time of the day or night, corresponding to the middle of the light and dark phases (respectively), to prevent any variation related to circadian gene expression. To avoid light-driven physiological responses, night samples were protected from light by aluminium foil and processed in low light conditions.

RNA extractions and purifications were performed using TRIzol reagent (Invitrogen), ethanol-isopropanol precipitation as described above, and RNA samples were resuspended in 50 μL RNAse-free H2O. RNA concentrations were measured by spectrophotometry using a NanoDrop One (Thermo Scientific), and DNA digestion was performed on 30 μL RNA samples using the RQ1 DNAse (Promega) following manufacturer’s instructions. After ethanol-isopropanol precipitation, DNA-digested RNA samples were resuspended in 30 μL RNAase-free water, and RNA integrity was assessed by electrophoresis on 1% agarose gel. Final RNA amounts ranged from 1 μg to 10 μg/sample, which were conserved at -80°C until library preparation.

#### Library preparation and RNAsequencing

Library preparation and RNAsequencing were performed at BGI Tech Solutions (Poland) using a DNBSEQ Eukaryotic Strand-specific Transcriptome Resequencing methodology. Briefly, mRNAs bearing polyA tail were enriched using oligo-dT beads, and were subsequently fragmented. First strand cDNA synthesis was performed using random primers, and second strand cDNA was synthesised with dUTP instead of dTTP. Double-stranded cDNA was subjected to end-repair and 3’ adenylated, before adapter ligation of the 3’ adenylated ends. PCR amplification using adapter primers was performed, and libraries were circularised after quality control. DNA nanoballs (DNB) were generated through library amplification by rolling circle replication, and sequenced using the DNBSEQ technology (37).

Raw sequencing data were filtered by the BGI Tech Solutions (Poland) using the SOAPnuke software. Briefly, adapters were removed from the sequences, and read with a length lower than 100 bp were discarded. Reads with polyX (A, T, G or C) sequences of more than 50 bp were also discarded, as well as reads with overall low quality (30% or more of the read with a quality value lower than 20). FASTQ files are available under the BioProject PRJNA1107376 accession number.

#### Read quality control and mapping

Quality control and mismatched base pair correction of filtered reads was performed by overlapping analysis using the –correction option of the fastp v0.24.0 tool (38).

For nuclear genes, genome index was generated using the STAR tool v2.7.11b, with default values (39). The NCBI RefSeq genome of *P. tricornutum* (GCF 000150955.2 ASM15095v2.fna) and its associated version 3 annotations (genomic.gtf) were used (5). Read mapping was performed using the same STAR tool and the previous genome index, with the –quantMode GeneCounts and – outSAMtype BAM SortedByCoordinate options. Unstranded read counts from all samples were combined in the ReadCount_Unstranded_grouped.csv file used for subsequent analyses.

For plastidial genes, the same procedure was followed using the *P. tricornutum* chloroplast genome (chromosome.chloroplast.fa) and a custom gtf annotation file generated from the complete chloroplast genome sequences (NC_008588.1), available in Dataset S5 (40).

#### Functional analysis from RNAseq data

Analyses were done with R (version 4.4.1). Differential gene expression was performed using DESeq2 v1.38.3 (41), which allows to test for differential expression based on a model using the negative binomial distribution, with default parameters. Comparison between control and *xatpa* strains was done with the “∼ type” design to only account for the effect due to the xATPA presence/absence, while the “∼ treatment + type + treatment:type” design was used for comparisons across culture conditions (treatments), to take into account the possible interaction between genotypes (*xatpa* or control strains) and culture conditions. Tables of log2 fold-changes and adjusted p-values are available in the Dataset S5. Genes with an absolute log2 fold-change above 2 and an adjusted p-value below 0.05 were considered to be differentially expressed. Regularised log2 counts were computed using the rlog function to allow variance stabilisation for small counts, and used for subsequent analyses.

Gene set enrichment analyses (GSEA) were performed to identify classes of genes over-represented in up or downregulated categories. Figures were produced using Gene Onthologies (GO) from genome wide annotation for Arabidopsis thaliana based on mapping using TAIR identifiers (org.At.tair.db v3.22.0 package with the clusterProfiler v4.6.2 package, without any gene filtration based on their log2FC or adjusted p-value, with default parameters (pvalueCutoff = 0.05). Functional enrichments observed were confirmed with two other approaches, with KEGG annotations (’pti’ kegg code) using the gseKEGG function of the clusterProfiler package, and with GO annotations using the topGO v2.50.0 package.

Weighted gene co-expression network analysis (WCGNA) was used to identified gene co-expression networks, and performed with the WGCNA v1.73 package (42). Computation was performed based on DESeq2 output, with the “∼ treatment + type + treatment:type” design, and using genes with at least 3 counts across the dataset and regularised log2 values. Modules were calculated using the blockwiseModules function, with a power of 11, for signed network, with a minimum and maximum module sizes set to 30 and 4000 respectively. Modules eigengenes were then computed and used to assess their Pearson correlation coefficients with experimental conditions, and GSEA analysis using KEGG annotation was performed with the clusterProfiler package as described above.

### Molecular properties of the xATPA protein in the F_1_ complex

#### Heterologous overexpression in ΔATP1 Saccharomyces cerevisiae yeast

The BY4741 *S. cerevisiae* strain knocked-out for the nucleus-encoded mitochondrial α subunit (Δ*ATP1*) (Euroscarf collection, Y03125). The complete genotype of this strain is MATa;his3D1; leu2D0; met15D0; ura3D0; atp1::KanMX4. The coding sequence of *S. cerevisiae ATP1* was amplified from genomic DNA, with its mitochondiral targeting sequences (MTS) (Dataset S3). For *P. tricornutum* genes, forward primer bearing the *ATP1* MTS was used to amplify *xATPA* and plastidial *ATPA* from genomic DNA, with *ATP1* MTS replacing *xATPA* plastidial targeting sequence (Dataset S3). Coding sequences were inserted by SLIC (sequence- and ligation-independent cloning) (43) in a yeast expression plasmid under the control of the strong GPD promoter, bearing the *ura3* selection gene (Dataset S3).

For each construct, 12 transgenic strains were selected on CSM-Ura 2% glucose. For spot assays, exponential cultures were prepared in liquid YPD, 30°C, shaking. Cells were pelleted, washed in liquid CSM-Ura 2% glucose and adjusted to DO=1. Serial dilution were performed in CSM-Ura 2% glucose, and 10 μL were spotted on solid CSM-Ura 2% glucose or CSM-Ura 2% glycerol (non-fermentable carbon source), and grown at 30°C.

#### Purification of double-tagged recombinant proteins

xATPA, plastidial ATPA and plastidial ATPB coding sequences were amplified from *P. tricornutum* genomic DNA using PrimeSTAR GXL DNA polymerase (Takara) and primers listed in (Dataset S3). Amplicons were inserted by restriction - ligation in IPTG-inducible expression plasmids between 5’-CBP/3’-6His (for ATPB) or 5’-TwinStrep/3’-6His (for xATPA and ATPA) purification tags (Dataset S3). The following protocol was adapted from (44).

Eukaryotic codon-optimised *E. coli* (Rosetta DE3 strain) were transformed by heat shock and selected on LB plates. Liquid seed cultures were grown overday (37°C, shaking) by inoculating a single transformed colony in 20 mL LB containing 100 μg/mL kanamycin. Seed cultures were then transferred for protein expression in 1 L Auto Induction LB (Formedium) containing 50 μg/mL of both kanamycin and chloramphenicol, and grown overnight at 28°C with 220 rpm shaking.

Buffer compositions are available in Dataset S4. Bacteria were harvested by centrifugation (15 min, 7 500 g), resuspended in 25 mL PBS, and centrifuged again (10 min, 6100 g). Pellets were resuspended in 25 mL lysis buffer and sonicated on ice (4 min, 30% amplitude, 1 s pulsation, 1 s pause, Brandson Digital Sonifier), resulting in the obtention of a clear solution. Cell fragments (P fraction) were pelleted by centrifugation (4°C, 25 min, 27 000 g), and 500 μL of cold His60 Superflow Resin slurry (Takara) pre-equilibrated in lysis buffer was added to the supernatant (soluble protein fraction). Tubes were left for 2h at 4°C on a rotating wheel to allow the binding of 6His-tagged proteins onto the resin. Beads were separated from the flowthrough (FT fraction) by gentle centrifugation (4°C, 2 min, 1 500 rmp), and washed with 10 mL cold lysis buffer. Beads were then washed with 750 μL cold wash buffer (W fraction), and eluted in 500 μL elution buffer (E fraction). Elution fractions were collected after gentle centrifugation (4°C, 1 min, 1 500 rmp), and a total of five elution steps were performed. P, FT, W and all E samples were examined on SDS-PAGE to identify fractions enriched in recombinant proteins.

Elution samples containing the protein of interest were pooled and placed in a 14 kD cut-off cellulose dialysis tube (Sigma Aldrich) and dialysed overnight in the appropriate interaction buffer under low stirring. Subsequent purification steps were performed in the same manner, starting on dialysed samples, using the relevant purification resins, wash and elution buffers. Strep-Tactin Sepharose (iba) was used for TwinStrep tag purification, and Calmodulin Affinity resin (Agilent) was used for CBP tag purification. Double-purified proteins were ultimately dialysed in storage buffer, aliquoted, flash-frozen in liquid N2 and conserved at -80°C.

Recombinant protein concentrations were assessed on SDS-PAGE (10% acrylamide) followed by Coomassie blue staining with BSA dilutions as references, and band intensities were analysed on ImageJ. Briefly, lanes of interest were selected with the gel analysis tool, and the area of the corresponding peak intensity was measured on the lane plot. Purified protein concentrations were then calculated using the standard curve built with diluted BSA samples.

#### In vitro interaction assay with purified proteins

The following protocol was adapted from (44). Interactions were performed in low protein binding tubes, using final concentrations of 0.1 μg/μL of each purified proteins, 0.4 μg/μL BSA and 2 mM dNTP (each), in storage buffer with a final volume adjusted to 50 μL. All components were carefully added to avoid denaturation, starting with purified proteins, and interaction mixes were incubated 30 min at 26°C to allow complex formation. 50 μL of the appropriate purification beads were pre-equilibrated in the appropriate interaction buffer containing 20 μg/mL BSA, resuspended in 500 μL of the same solution, and added gently to the interaction tubes. Tubes were left 1h30 at 4°C on a rotating wheel to allow the binding of tagged-proteins onto the purification resin.

Beads were pelleted by gentle centrifugation (4°C, 5 min, 1 500 rmp), and washed with 500 μL of cold interaction buffer. After two washes, elutions were performed by adding 20 μL of the appropriate elution buffer and by incubating the tubes 10 min at 30°C. Elution fractions were collected by centrifugation (2 min, 2000 g), and concentrated by evaporation by incubating open tubes at 95°C. Sample contents were visualised by SDS-PAGE (10% acrylamide) followed by silver staining.

#### In vitro affinity measurement by Surface Plasmon Resonance

Measurements were performed on a Bioacore 3000 (GE Healthcare). CM5 chips (Cytiva) were first activated by 0.4 M EDC and 0.1 M NHS, and purified CBP-ATPB-6His proteins were then covalently immobilised on the surface by injection of 50 nM protein in 10 mMsodium acetate pH = 4.5 (Cytiva), until a value of 4000 Ru. The surface was then deactivated with ethanolamine HCl pH = 8.4 (Cytiva).

Interaction assays were performed in HBS-EP pH = 7.4 buffer (Cytiva), used as running and dilution buffer for partner proteins. The flow was set up at 5 μL/min, complex association occurred during 5 min upon partner protein injection, and dissociation was monitored upon running buffer injection. Chip regeneration was performed for a 30 s duration by injection of 15 μL of 10 mM Glycine-HCl pH = 2.5, allowing total complex disassembly, followed by a 5 min equilibration step with 5 μL/min running buffer.

Measurements were replicated on two independent CM5 chips. Interaction kinetics were established for 3 to 300 nM TwinStrep-xATPA-6His and 0.4 to 275 nM for TwinStrep-ATPA-6His. Data analysis was performed with the constructor BIAeval software, using Langmuir 1:1 and steady state affinity models.

#### Pull-down on P. tricornutum whole cell lysate

100 mL *Phaeodactylum tricornutum* cultures in exponential phase (1∼5x10^6^ cells/mL) grown in standard conditions were harvested by centrifugation in the middle of the light phase, and resuspended in 500 μL TwinStrep interaction buffer. Cells were pelleted in low protein binding tubes, and frozen in liquid N2. Frozen pellets were crushed on ice with a tube pestle until completely thawn, and 900 μL of TwinStrep interaction buffer supplemented with 100 μL 10X cOmplete protease inhibitor (Sigma Aldrich) were added. Cells were then sonicated on ice (10 min, 20% amplitude, 30 s pulsation, 1 min pause, Bioblock Scientific Vibracell 75041), resulting in clear solutions. Cell fragments were pelleted by centrifugation (4°C, 30 min, 20 000 g), and BCA quantification (Millipore) was used to estimate supernatant protein concentrations.

Interactions were performed in low protein binding tubes (Eppendorf), using 6 μg of whole cell lysate proteins, 150 ng of purified bait protein, with a final volume adjusted to 500 μL using TwinStrep interaction buffer. Tubes were left overnight at 4°C on a rotating wheel to allow the formation of protein complexes. 50 μL of Strep-Tactin Sepharose (iba) was pre-equilibrated in TwinStrep interaction buffer and added to the interaction tubes. Tubes were left 1h at 4°C on a rotating wheel to allow the binding of tagged-proteins onto the purification resin.

Beads were pelleted by gentle centrifugation (4°C, 5 min, 1 500 rmp), and washed with 500 μL cold TwinStrep interaction buffer. After a total of three washes, elutions were performed by adding 32 μL of TwinStrep elution buffer and by incubating the tubes 15 min at 30°C. Elution fractions were collected by centrifugation (2 min, 2000 g), and conserved at -80°C.

#### Protein identification by mass spectrometry

Protein samples were thawed by incubation in an ultrasonic bath for 5 min, and then heated to 90°C for 5 min before being deposited on Criterion gels (10% acrylamide for separation, 4% for concentration). Migration was performed with a constant current of 20 mA over approximately 3 mm in the separation gel. After migration, gels were fixed and then stained with silver nitrate.

Lanes were cut and decolourised prior to tryptic digestion. Disulfide bridges were reduced by incubating the gel pieces in a 10 mM dithiothreitol (DTT) solution for 30 min at 56°C. Cysteine alkylation was performed by adding a 50 mM iodoacetamide (IAA) solution, followed by incubation for 30 min in the dark. Gel pieces were washed with a 50 mM ammonium bicarbonate solution and then dehydrated with acetonitrile. Tryptic digestion was performed overnight at 37°C with 300 ng of trypsin (mass spec sequencing grade Trypsin, Promega) per sample. Resulting peptides were extracted with two incubations in a 60% acetonitrile/0.1% TFA solution in an ultrasonic bath for 10 min. Samples were dried completely in a Speed-Vac, then resuspended in 20 μL of a 0.1% formic acid/2% acetonitrile solution and desalted by StageTips (Affinisep T2 tips) with the DigestPro MSI robot (Intavis).

Samples were processed using the nanoLC-MS system nanoElute II coupled with TimsTOF HT mass spectrometer (Bruker). LC separation was done using a 30-minute gradient of acetonitrile (5-16%/22-27%/27-95%) at 220 nL/min on a C18 reverse phase column (Aurora3 Ultimate – IonOpticks, length 25 cm, diameter 75 μm, particles of 1.7 μm, pores of 12 nm), coupled with a 0.22 μm pre-column filter (EXP2, Optimise Technologies). Data were acquired using Data Independent Acquisition (DDA), with ion mobility activated (TIMS, mobility range 0.7–1.3 1/K0) and CID fragmentation.

LC-MS/MS analysis was performed using MaxQuant software version 2.7.0.0, with parameters set to instrument = Bruker TIMS; first search error tolerance = 20 ppm, main search error tolerance = 10 ppm; enzyme = trypsin/P, specific, max missed cleavages = 3; modifications = acetyl (N-term) and oxidation (M) as variable modifications, carbamidomethylation of cysteine as a fixed modification in a maximum of 3 modifications per identified peptide; quantification = LFQ (minRatio 1, ALL peptides); normalization type = NONE. The *Phaeodactylum tricornutum* UniProt reference proteome (downloaded on 26/06/2025) was used for peptide identification, with TwinStrep-ATPA-6His and TwinStrep-xATPA-6His sequences added, as well as the DB Contaminants MaxQuant list. The mass spectrometry proteomics data are accessible via the ProteomeXchange Consortium with the dataset identifier PXD080074.

Identification and abundance results were analysed the following R packages : FactoMineR, factoextra, dplyr, tidyr, reshape2, ggplot2, ggrepel, ggVennDiagram and ggvenn. Proteins were renamed and assigned a subcellular localisation based on their UniProt annotations (accessed 22/08/2025). Due to the small number of identified proteins, their low abundances and the lack of a normalisation reference, MS/MS count values (discrete) were used rather than LFQ intensities (continuous). Protein counts are available in Dataset S5.

## Supplementary Figures

**Fig. S1.**
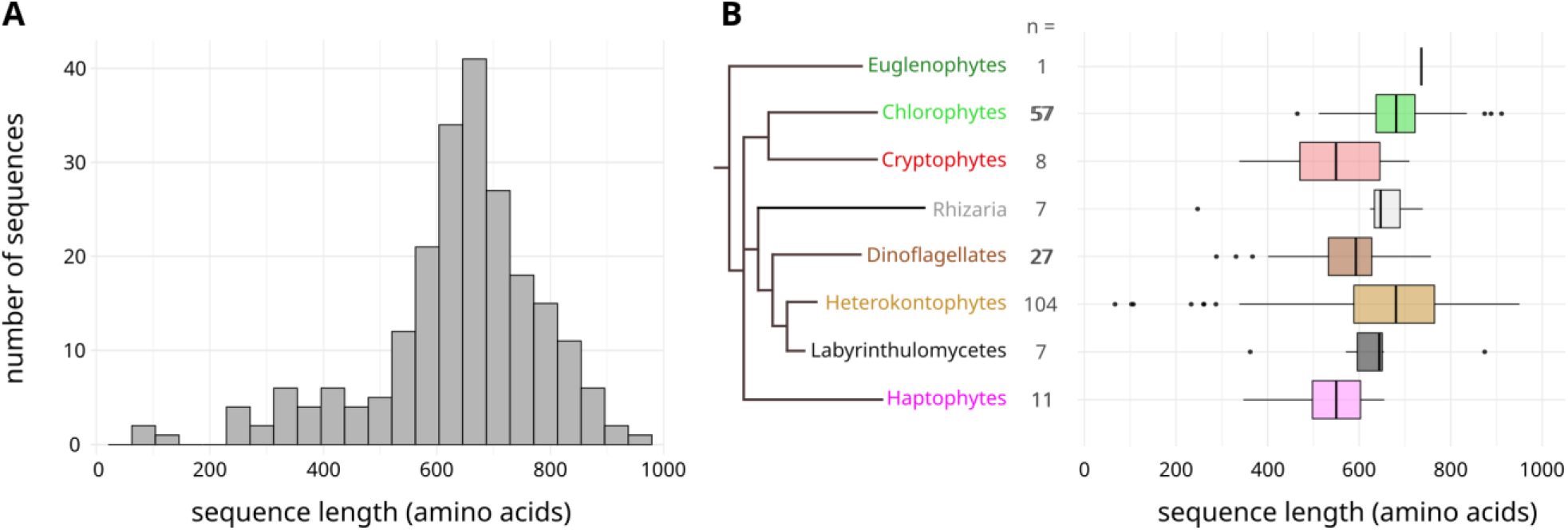
Length of xATPA in Eukaryotes. A. Sequence length of xATPA proteins in our dataset. B. Sequence length for each taxon. Boxplots show the length distribution of xATPA in the corresponding taxon. The number of xATPA sequences in each taxon is shown in the middle column.

**Fig. S2.**
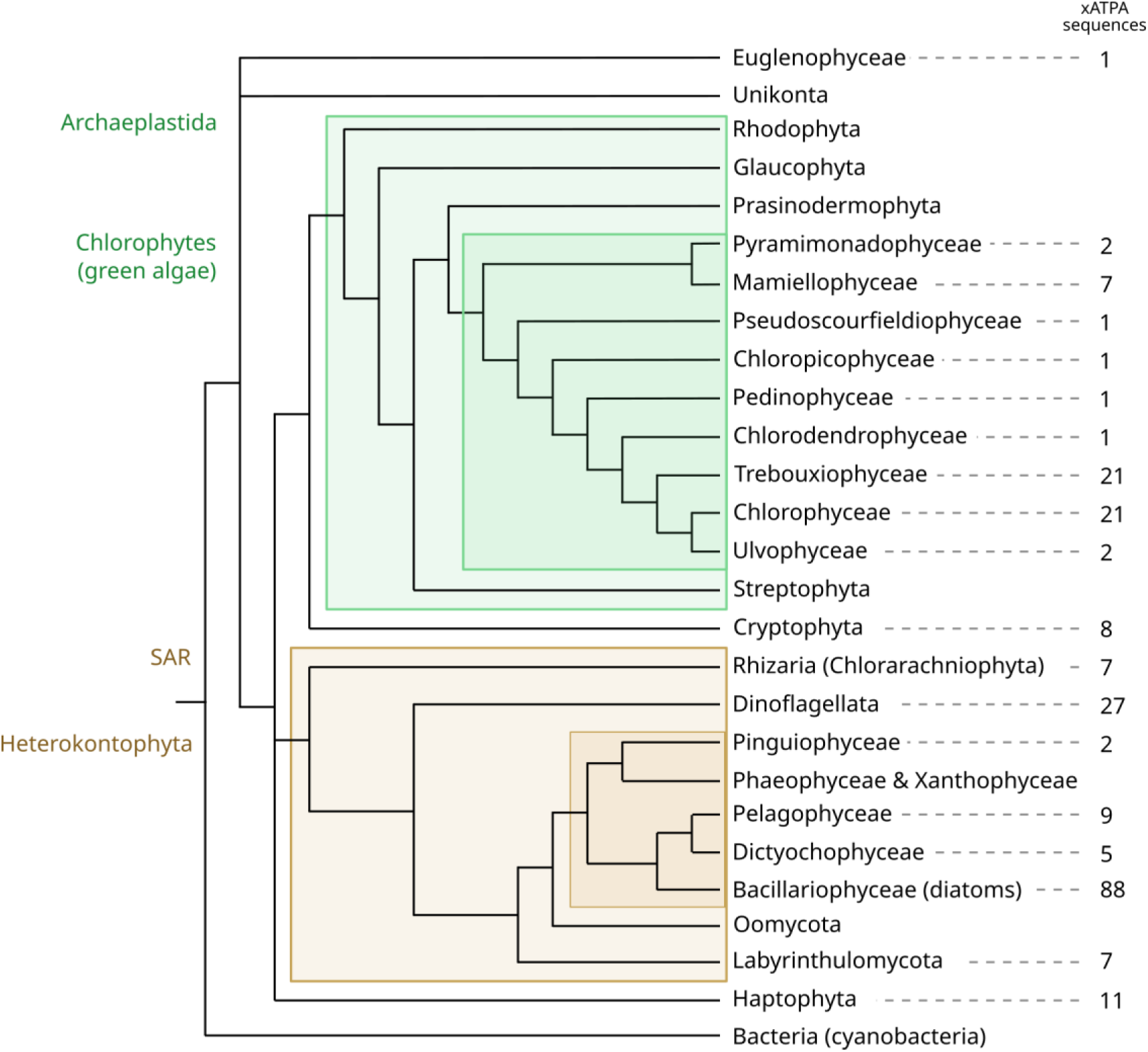
Distribution of the 222 xATPA homologues on a simplified Tree of Life. xATPA sequences were mined from more than 600 species, encompassing the whole diversity of the tree of eukaryotes, as well as Bacteria and Archaea. Our custom dataset is composed of both genomes and transcriptomes (Dataset S1), and has been complemented with more detailed searches on NCBI (accessed on 02/09/2025). The column indicates the number of xATPA sequences found in each group. Taxa not represented indicates an absence of xATPA detection

**Fig. S3.**
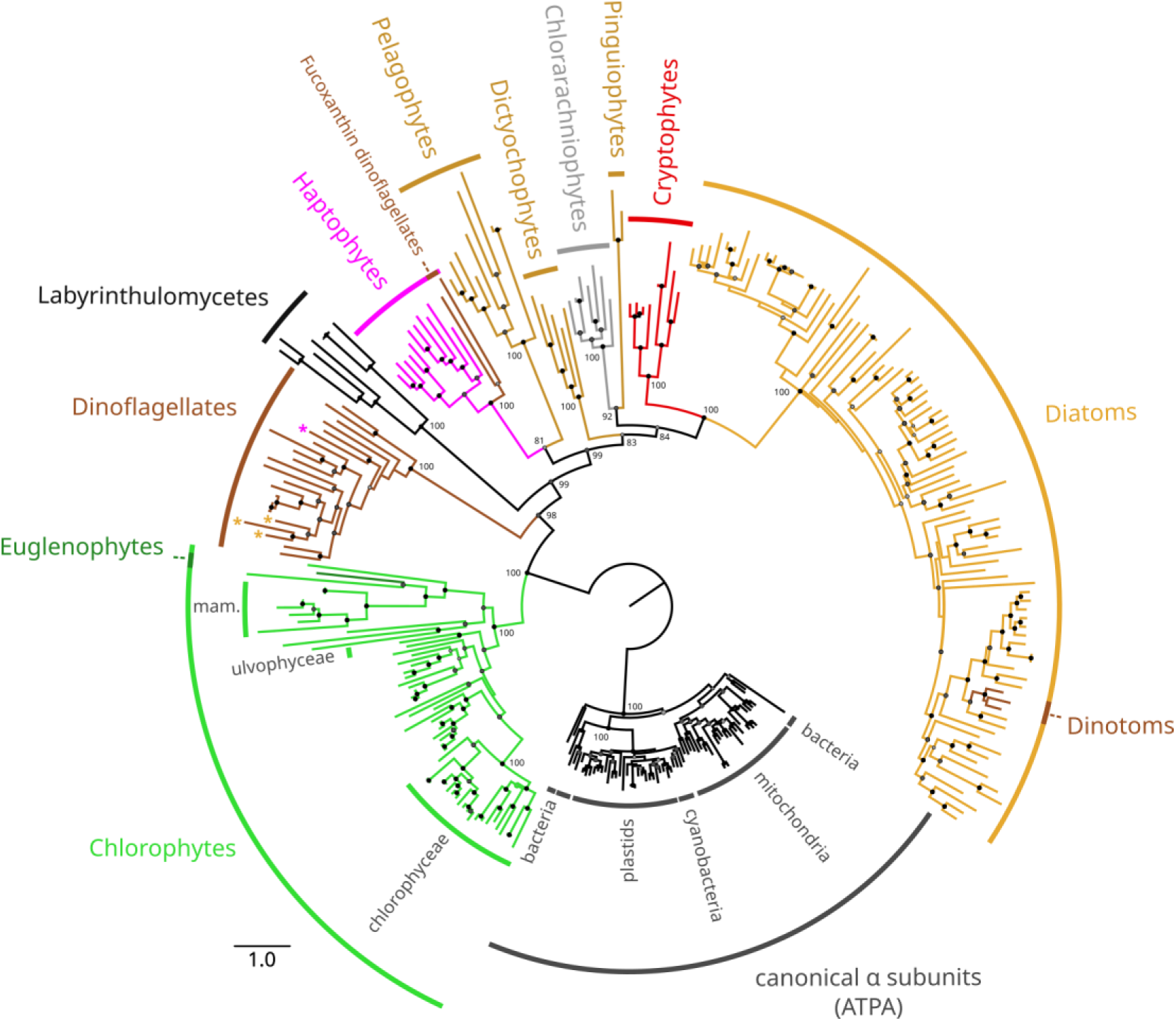
Phylogenetic tree of full xATPA proteins. Tree was built using IQ-TREE3 using the best substitution model Q.PFAM+F+R10. Canonical ATPA subunits from diverse origins were used as the outgroup. Branches are coloured based on their taxonomy, the scale bar represents the average amino acid mutation rate, and bootstrap branch supports higher than 75 are represented at each node and coloured based on their value: 100 = black, >90 = dark grey, >75 = light grey. Bootstrap values are shown on important nodes. In dinoflagellates, stars indicate species that have two xATPA copies, and their colours correspond to the taxa in which the second copy is found. A similar topology is obtained when canonical ATPB sequences are integrated to the analysis.

**Fig. S4.**
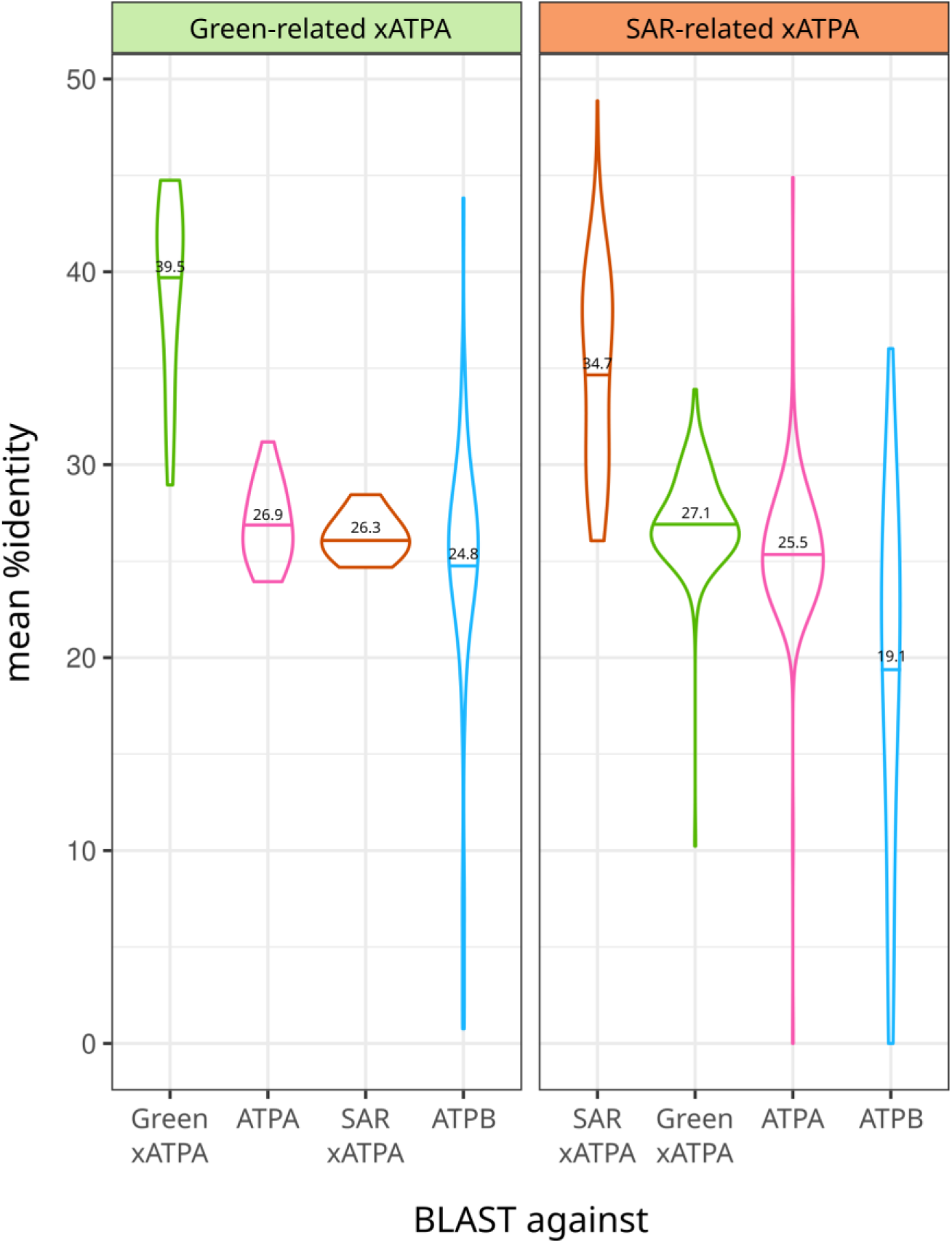
Sequence divergence from internal BLASTp comparison. An internal BLASTp was performed using the 222 xATPA sequences as queries against a database made of all xATPA and canonical ATPA (n=68) and ATPB (n=74) sequences. xATPA were classified as green-related (n=58) and SAR-related (n=164) based on their position in xATPA phylogenetic tree, and the mean percentage of identity to each category was computed. Violin plots visually confirm the distinction between green- and SAR-related xATPA. Mean is indicated by a bar and with its value. When considering all xATPA as a whole, mean sequence identity is significantly higher with ATPA (25.9%) than with ATPB (20.6%), based on a two-tailed paired t-test (p = 1.9 x 10^−16^, df = 221).

**Fig. S5.**
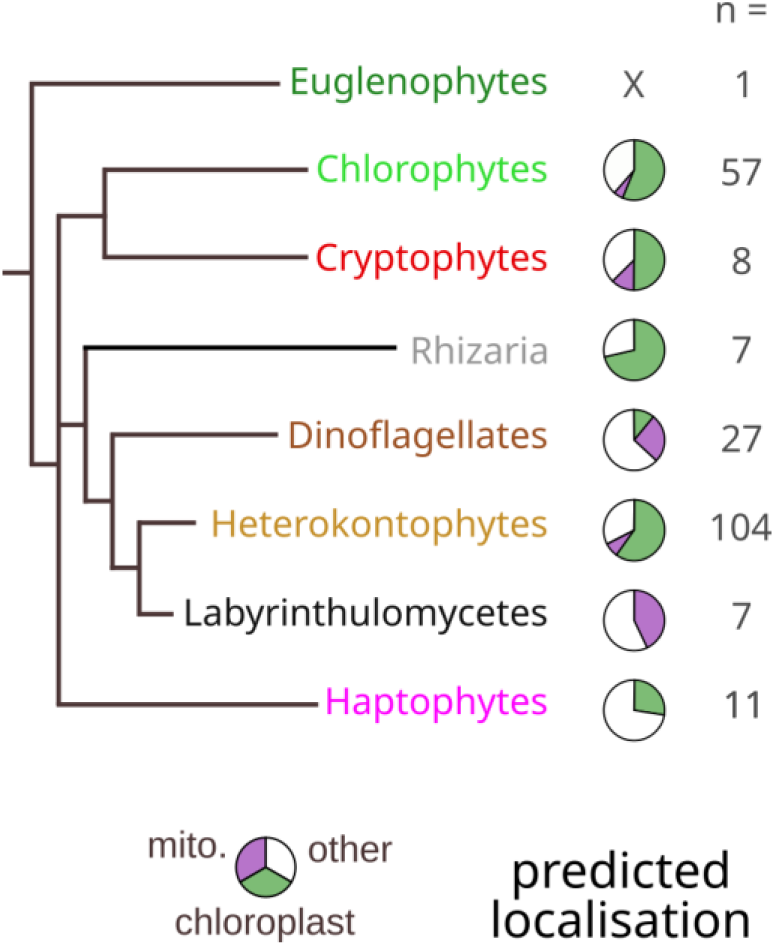
Predicted intracellular localisation of xATPA in eukaryotes. Predicted localisation for each taxon. Pie charts represent the proportion of subcellular targeting based on DeepLoc2 prediction in the corresponding taxon. The *Euglena gracilis* prediction is not shown, since this protein is already known to be plastidial (45). The number of xATPA sequences in each taxon is shown in the right column.

**Fig. S6.**
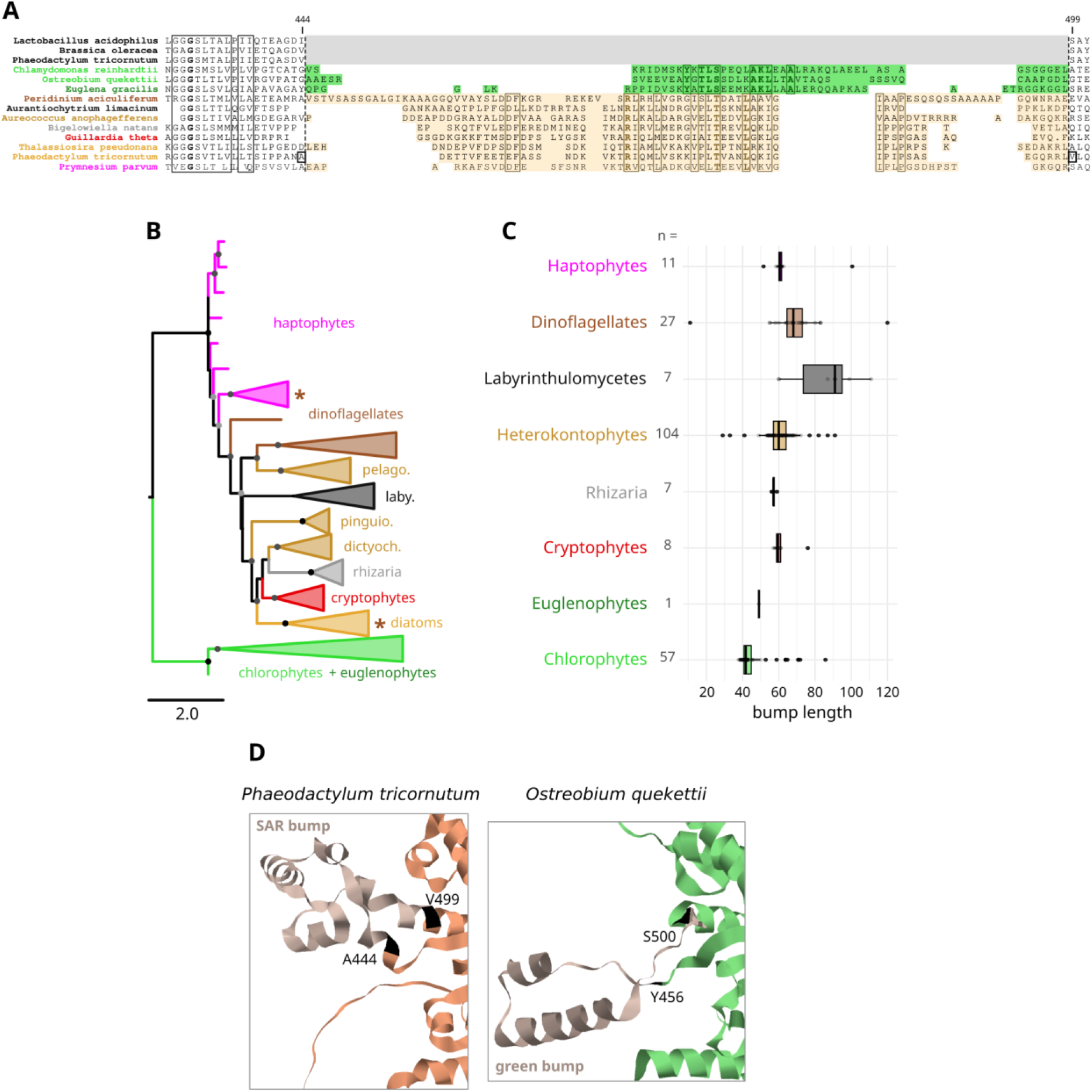
Green and SAR-derived xATPA bear different bump domains. A: Alignment of selected ATPA and xATPA around the bump domain, visualised using the ES-ript v3.2 online tool (Robert and Gouet, 2014). Bump domains are delimited by the dashed lines. The grey area shows its absences from ATPA, green and orange areas highlight the bump domain of green and SAR-derived xATPA, respectively. Frames indicate conserved sequences, and identical amino acids are presented in bold. Coordinates shown on top correspond to the amino acid position in the *P. tricornutum* xATPA sequence. B: Phylogenetic tree of the bump region. The tree was built with IQ-TREE3 using the best substitution model Q.PFAM+F+R5. Branches are coloured based on their taxonomy, scale bar represents the average amino acid mutation rate, the bootstrap branch supports higher than 75 are represented at each node and coloured based on their value: 100 = black, >90 = dark grey, >75 = light grey. Stars indicate clades including a species from a different group (endosymbiotic host), and are coloured correspond to this group. C: Bump domain length distribution in each taxon, considering frontiers defined by the dashed lines in A. D: Close-up on the bump region of predicted structures from the diatom *P. tricornutum* xATPA (orange, UniProt : B7FRE6, pLDDT = 76%) and the green alga *O. quekerttii* (pTM = 0.77). Bump domains are coloured in grey, flanking amino acids are shown in black and are numbered based on their position in each protein sequence

**Fig. S7.**
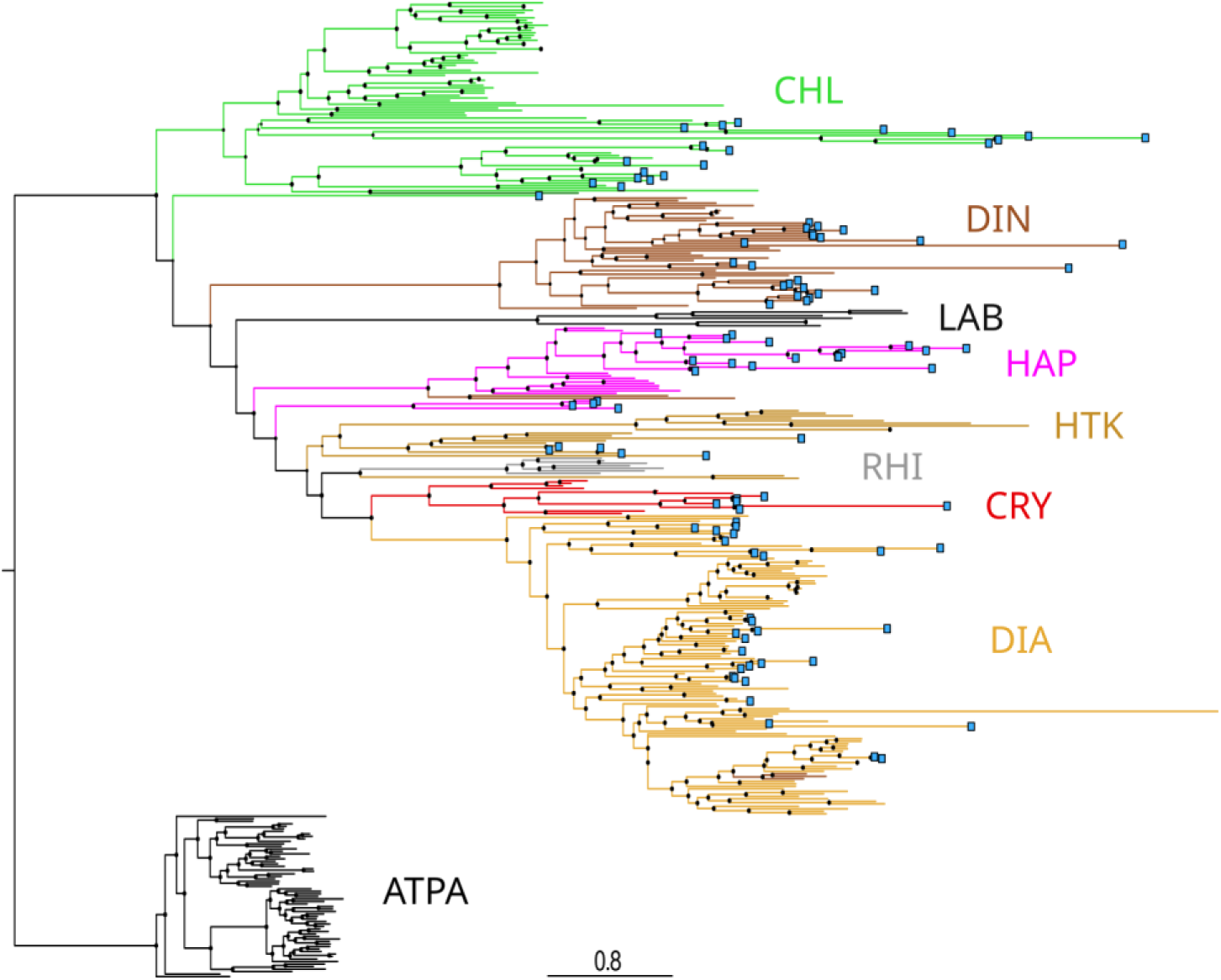
Phylogeny of environmental xATPA sequences. xATPA sequences were mined from the MATOU-v1.5 *Tara* Oceans dataset, and a phylogenetic tree of environmental xATPA unigenes was built based on a topology constrained by the ATP synthase domain phylogeny (Fig 1A) using IQ-TREE 3 with the substitution model Q.PFAM+F+R10. Environmental sequences are indicated with a blue squared tip. Branches are coloured based on their taxonomy, scale bar represents the average amino acid mutation rate, node sizes indicate bootstrap branch support. CHL : chlorophytes, DIN : dinoflagellates, LAB : labyrinthulomycetes, HAP : haptophytes, HTK : non-diatom heterokontophytes, RHI : rhizarians, CRY : cryptophytes, DIA : diatoms. Environmental sequences were taxonomically assigned based on their position in this tree.

**Fig. S8.**
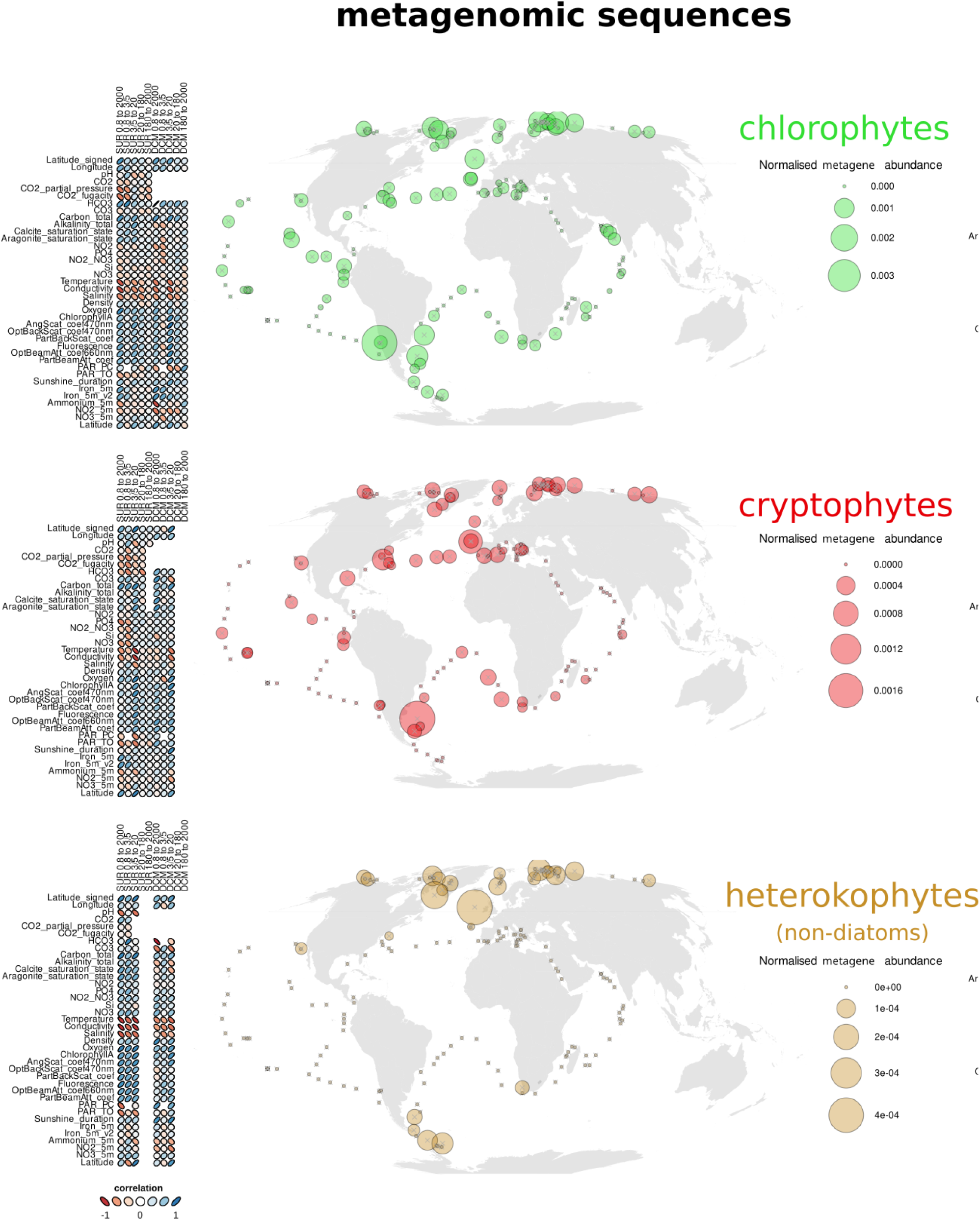
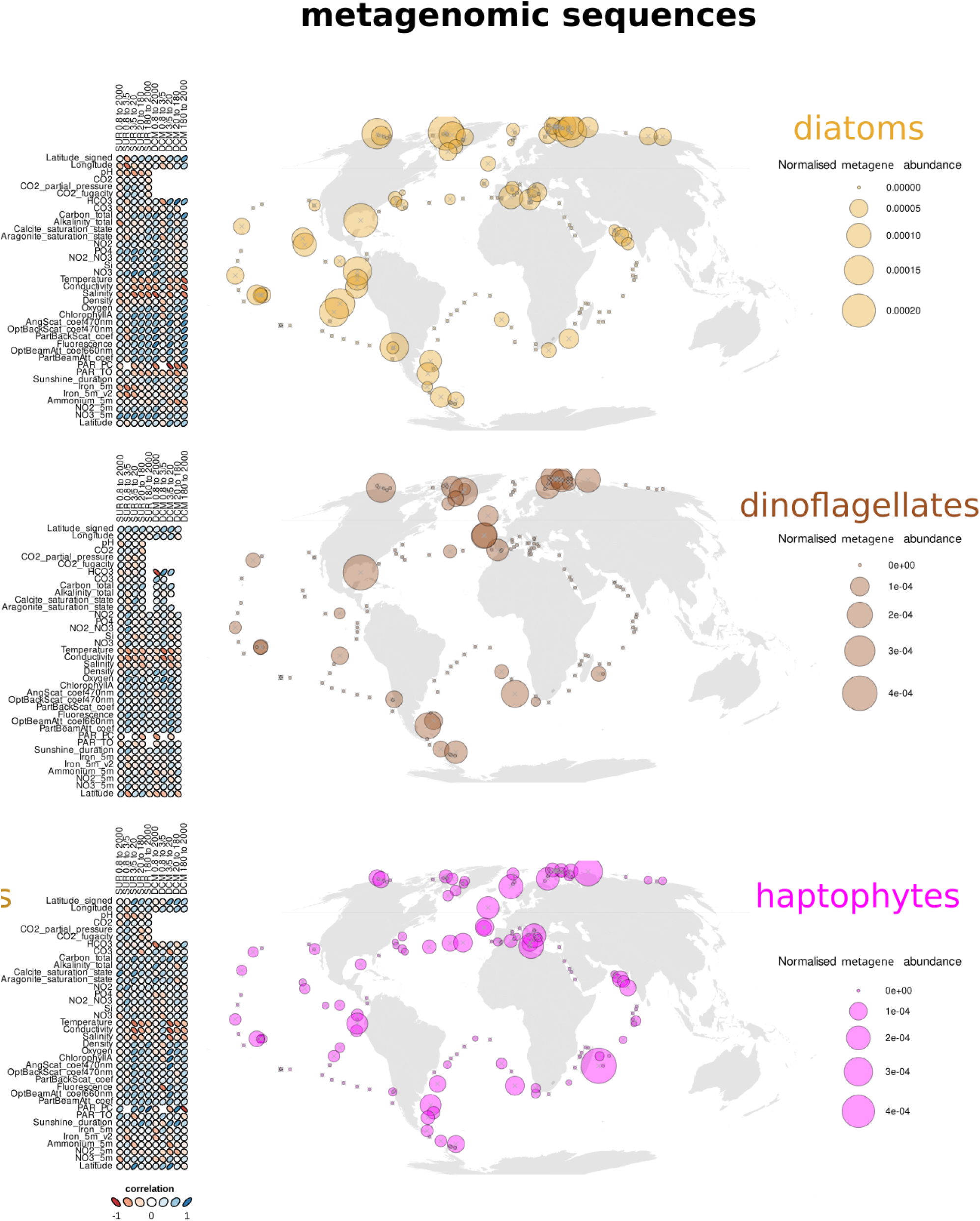
*xATPA* metagenomic distribution and correlation with ocean parameters. *xATPA* metagenomic abundances were normalised for each taxa based on the total metagenomic abundances of the corresponding taxa in each sample. Taxon is indicated next to the plot. Left: Spearman’s rank correlations between *xATPA* abundances from surface or DCM samples and environmental parameters. Each column corresponds to a different size fraction, indicated at the bottom. Correlation coefficients are represented by both ellipse shapes and colours (red = negative, blue = positive). Right: Distribution and abundances of *xATPA* metagenes in *Tara* Oceans sampling stations. Circle size represents *xATPA* abundances within a station, which is here the sum of normalised *xATPA* abundances from each sample of the same station, across all size fractions and depths.

**Fig. S9.**
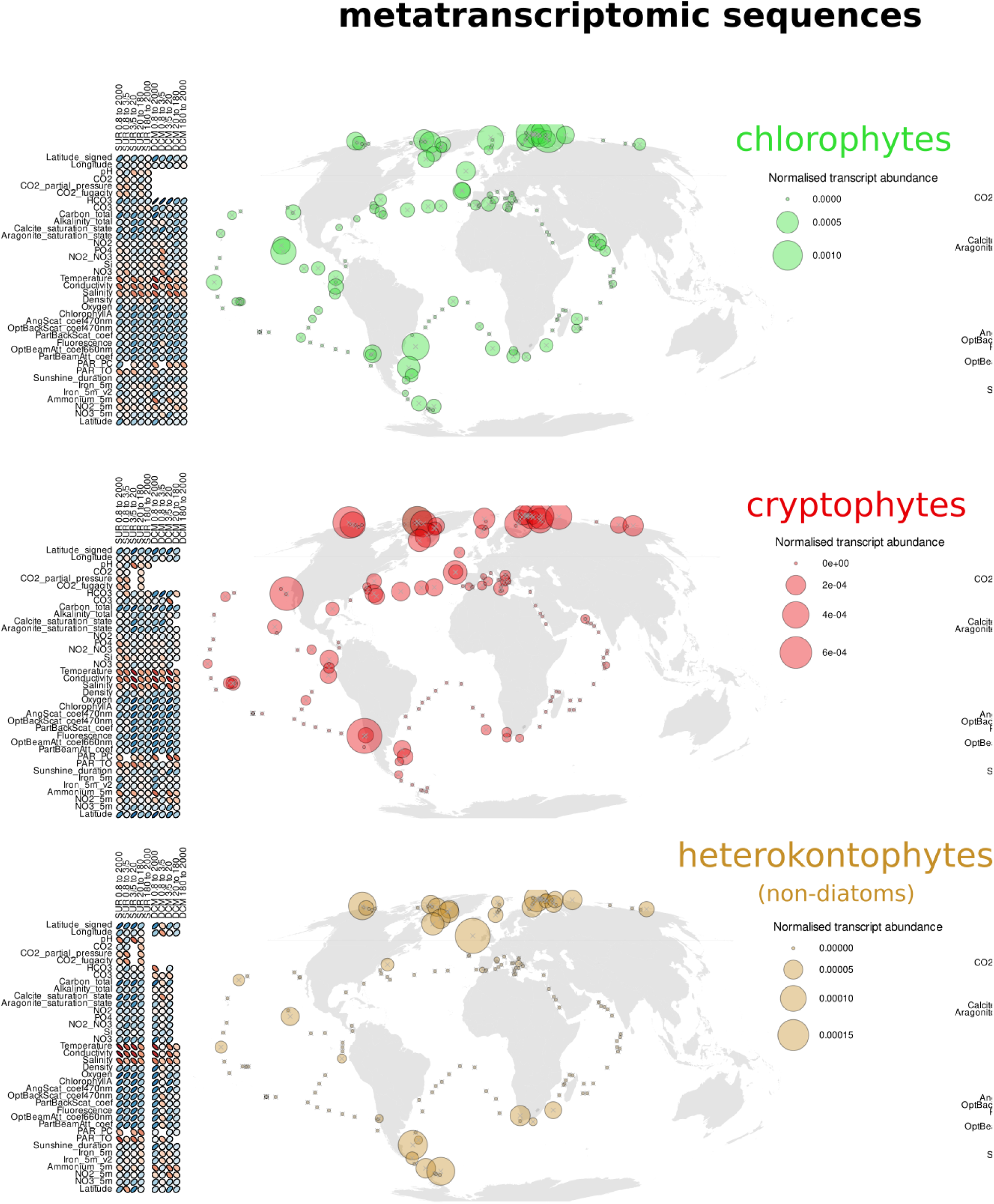
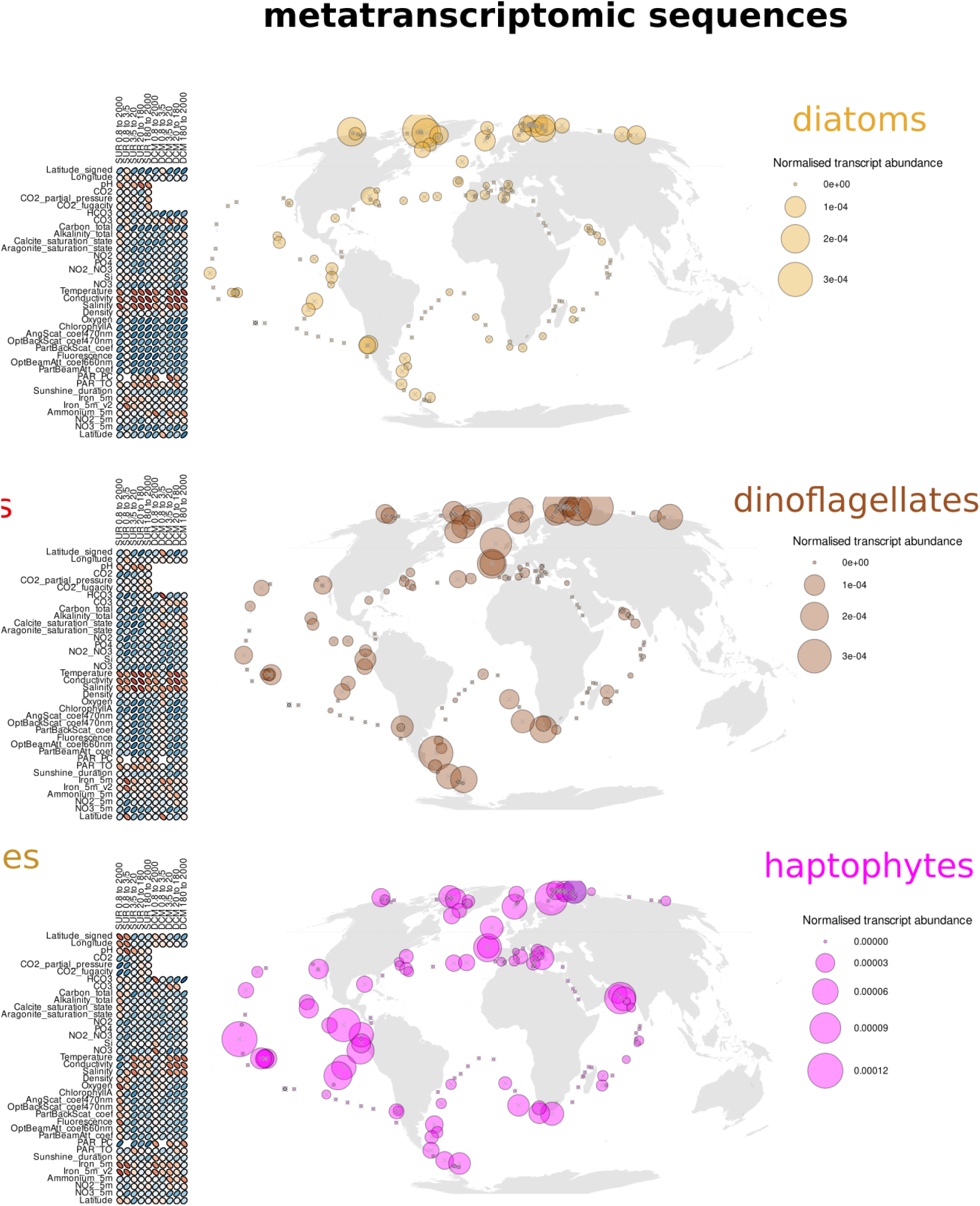
*xATPA* metatranscriptomic distribution and correlation with ocean parameters. *xATPA* metatranscriptomic abundances were normalised for each taxa based on the total metatranscriptomic abundances of the corresponding taxa in each sample. Taxon is indicated next to the plot. Left: Spearman’s rank correlations between *xATPA* abundances from surface or DCM samples and environmental parameters. Each column corresponds to a different size fraction, indicated at the bottom. Correlation coefficients are represented by both ellipse shapes and colours (red = negative, blue = positive). Right: Distribution and abundances of *xATPA* metatranscripts in *Tara* Oceans sampling stations. Circle size represents *xATPA* abundances within a station, which is here the sum of normalised *xATPA* abundances from each sample of the same station, across all size fractions and depths.

**Fig. S10.**
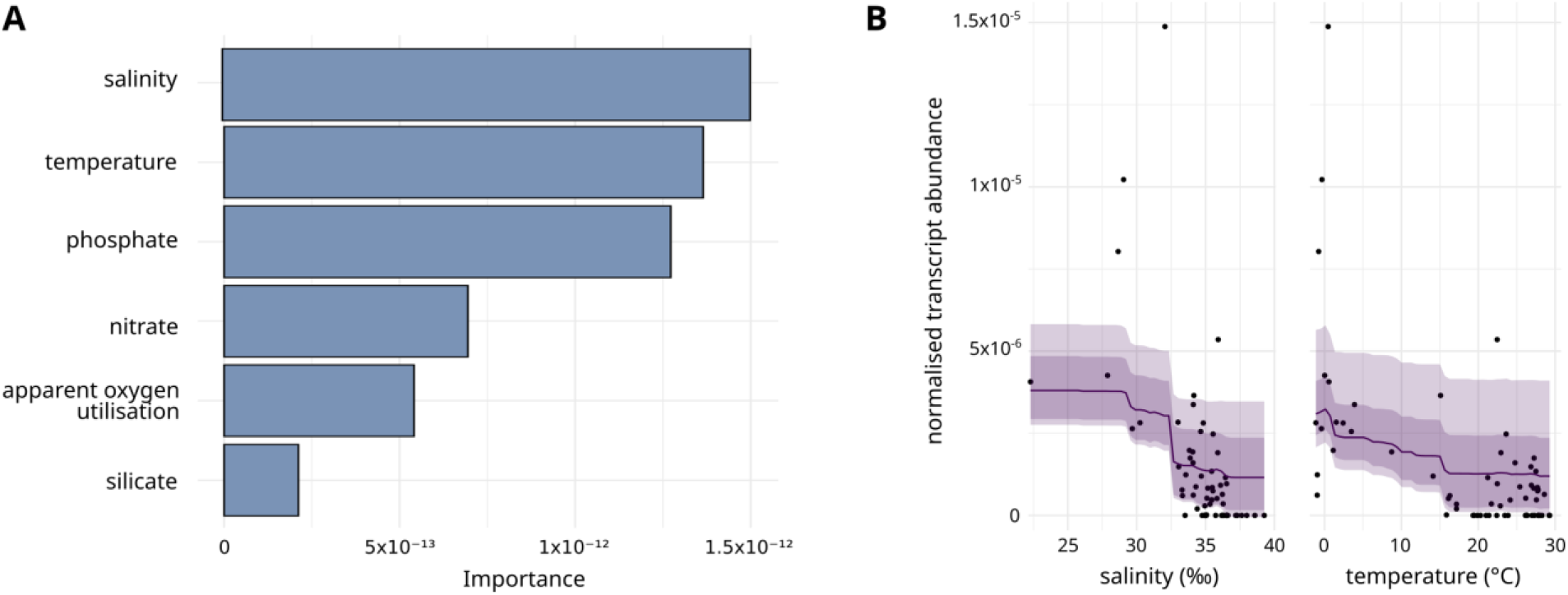
Environmental predictors of *xATPA* metatranscript abundances in diatoms at the ocean surface. Normalised metatranscript abundances used are the log10-transformed mean values calculated within each station in surface samples. These data corresponds to the random forests regression model used to predict abundances in Fig 3A (R^2^ = 0.82), based on measured *xATPA* abundance from *Tara* Oceans and global oceanic environmental variables from the World Ocean Atlas 2018. A: Importance of environmental predictors in the model. B: Partial dependency plots of the top two predictors. Measured values are represented as black dots. Abundance estimation means are represented by the solid lines, its standard deviations are represented by the dark envelope, and 5% and 95% centiles by the light enveloppe.

**Fig. S11.**
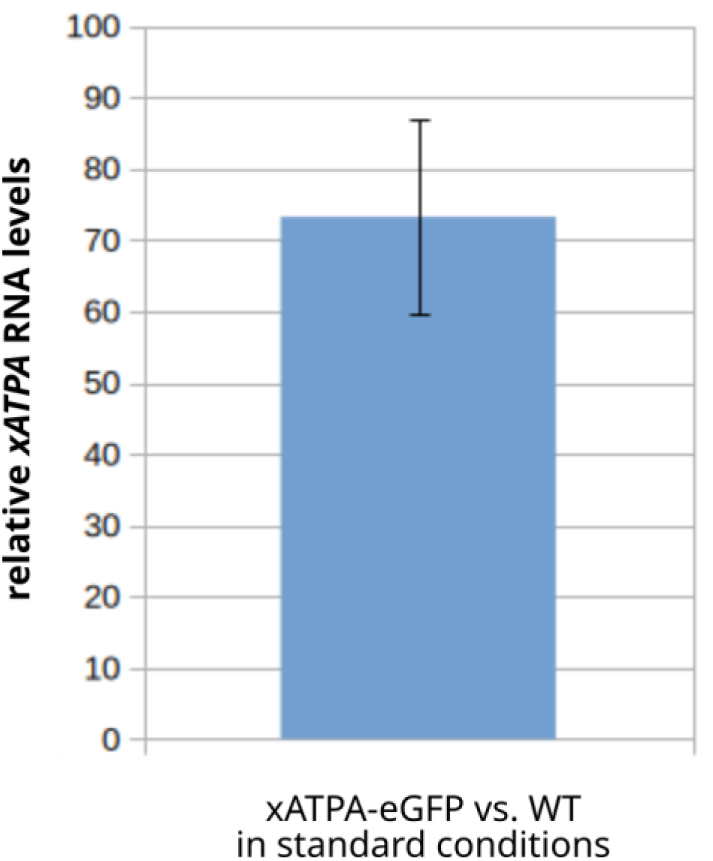
Overexpression of the xATPA-eGFP transgene in standard conditions at day mid-point. RT-qPCR was performed on samples from xATPA-eGFP and WT strains in exponential phase harvested at day mid-point, using primers targeting two different locii of the *xATPA* gene and the housekeeping RPS and TBP genes (36). Relative xATPA expression between xATPA-eGFP and WT strains was computed from four technical replicates (mean=73, sd=14, n=4). For a one-tailed (“greater”) Wilcoxon test with a null-hypothesis of mean=1 (ie no differerence in relative expression), the p-value is 0.0625. Hence, we consider reasonable to conclude that xATPA is overexpressed in the xATPA-eGFP strain.

**Fig. S12.**
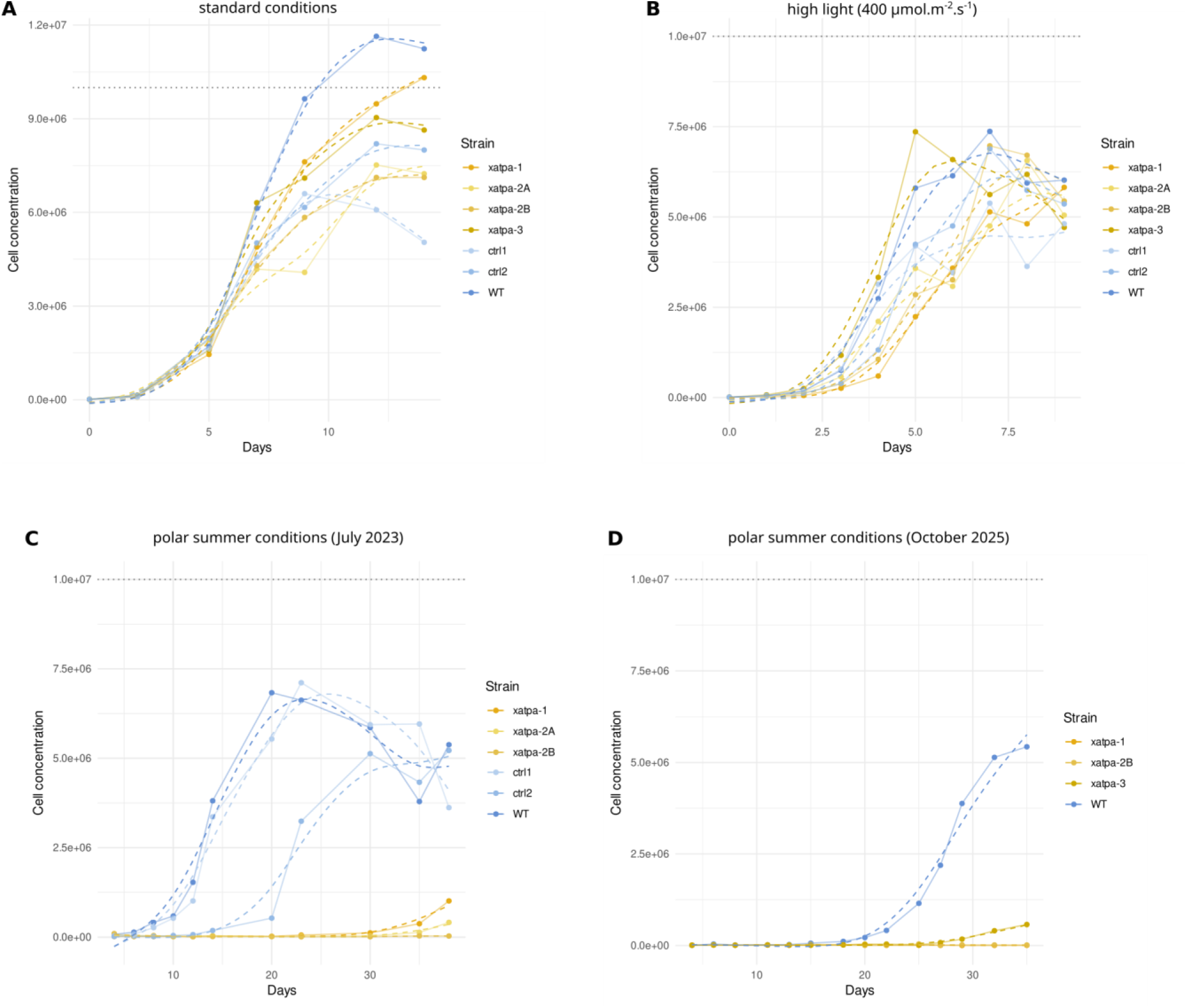
Individual growth curves in key environmental conditions tested. Cell concentration was followed during the growth, at a constant hour (day mid-point). Measured values are represented by points and solid lines, fitting curves (GAM) are represented by dashed lines, and colours correspond to the different strains. WT = wild type, *xatpa* = *xATPA* KO strains, *ctrl* = control strains (transformed with zeocine resistance and human Cas9 plasmids but without sgRNA). A horizontal dotted line indicate the concentration of 1x10^7^ cells/mL, allowing the comparison between all diagrams. A: standard condition, B: high light condition (12L:12D), C: polar summer conditions (July 2023), D: polar summer conditions (October 2025).

**Fig. S13.**
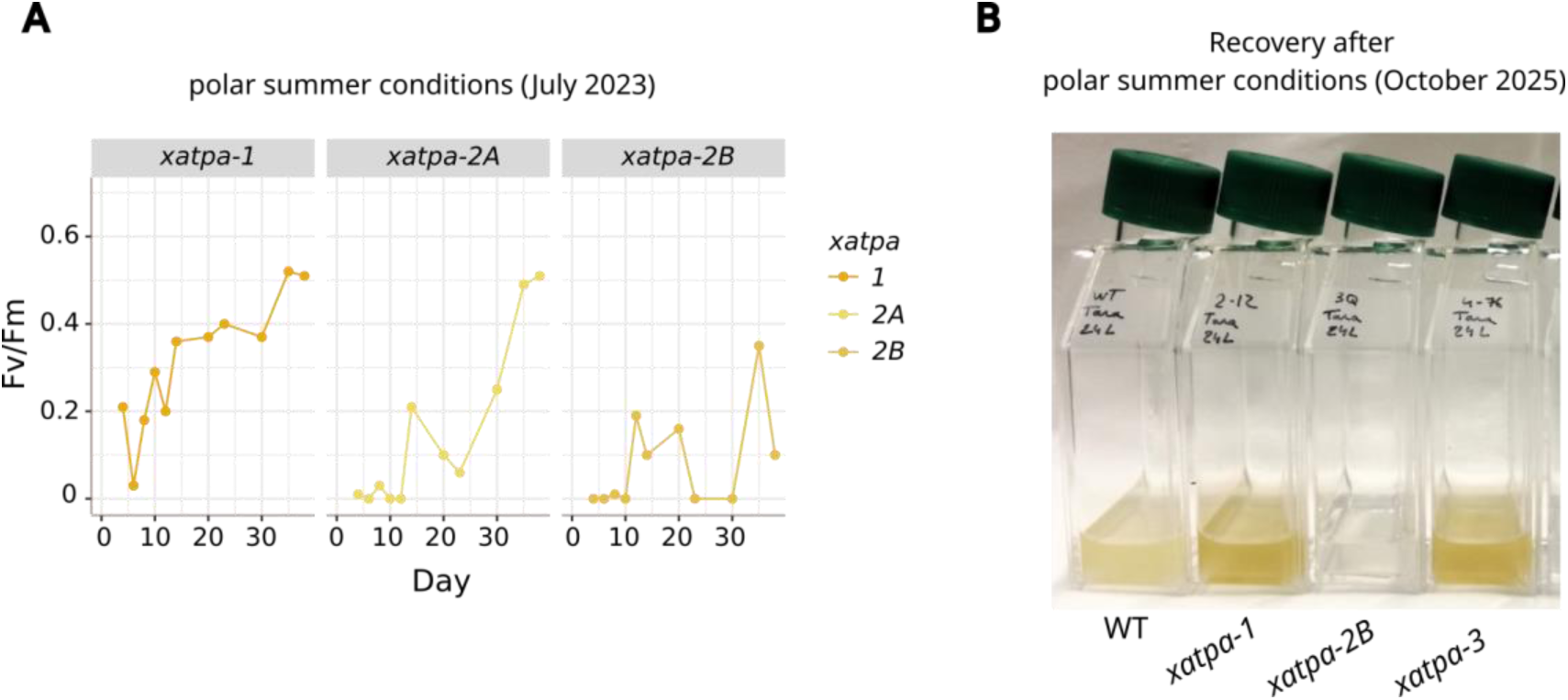
Viability of *xATPA* KO strains during and after culture in polar summer conditions. Growth curves were performed in polar summer conditions for a month, in two independent experiments. A: Fv/Fm of *xatpa* cultures was measured during the July 2023 growth curves in polar summer conditions. Photosynthesis efficiencies do not remain null, and cells can be considered viable. However, Fv/Fm low values suggest that cells experience cellular stresses. B: after the October 2025 growth curves in polar summer conditions, cultures were placed in standard conditions for a month to allow cells to recover. All *xatpa* strains resumed their growth, although the *xatpa*-2B strain showed lower cell density.

**Fig. S14.**
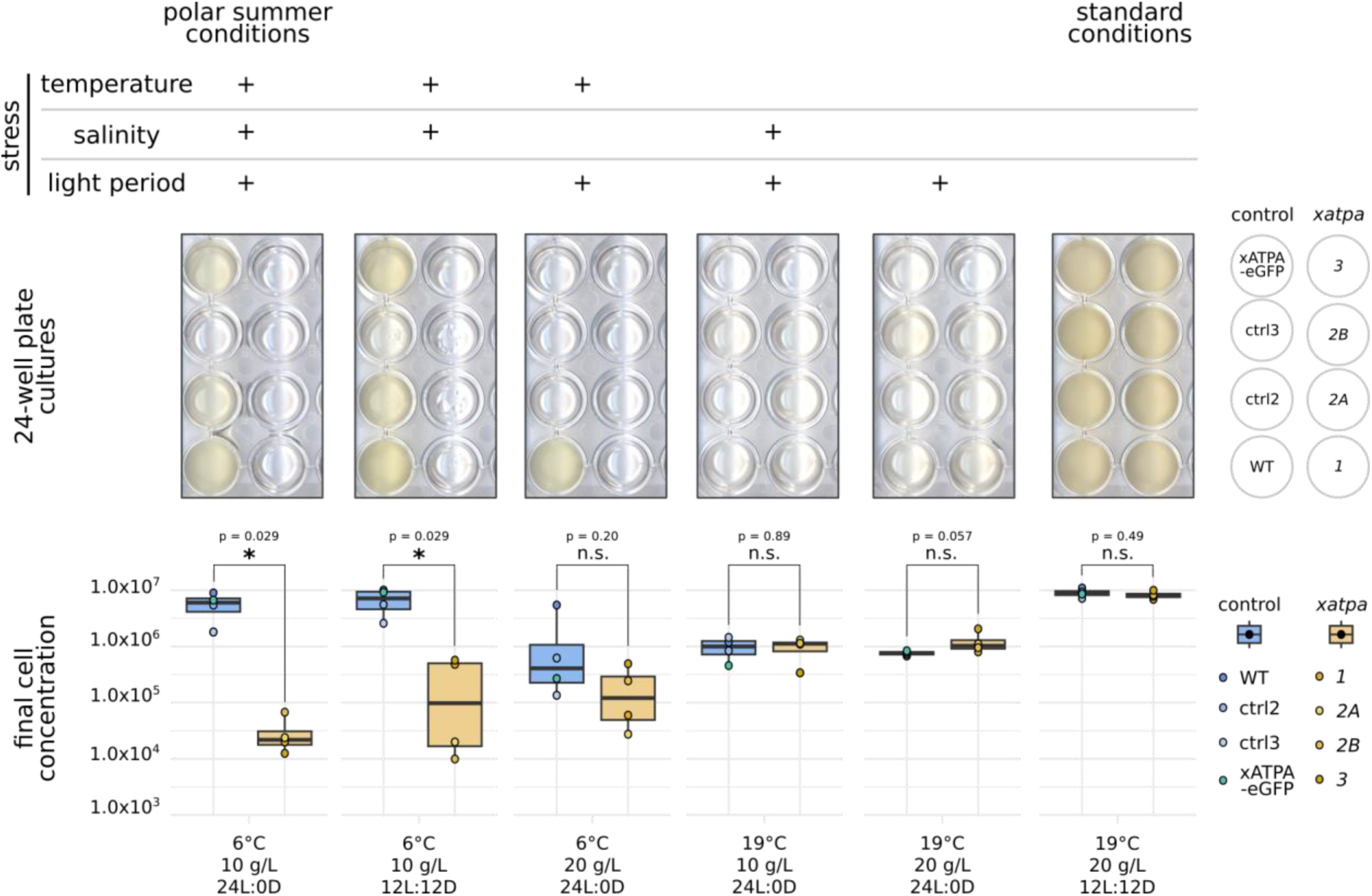
Growth abilities of *xatpa* strains in different combinations of polar summer-related stresses. Top: growth assays in 24-well plates, and Bottom: corresponding final cell concentrations. Two-tailed Wilcoxon tests were performed to assess the significance of mean differences (*: p<0.05, **: p<0.01, ns : non significant). For each growth condition, combinations of polar summer-related stresses and exact parameter values are indicated at the top and bottom of each picture, respectively. Left border: polar summer conditions. Right border: standard conditions

**Fig. S15.**
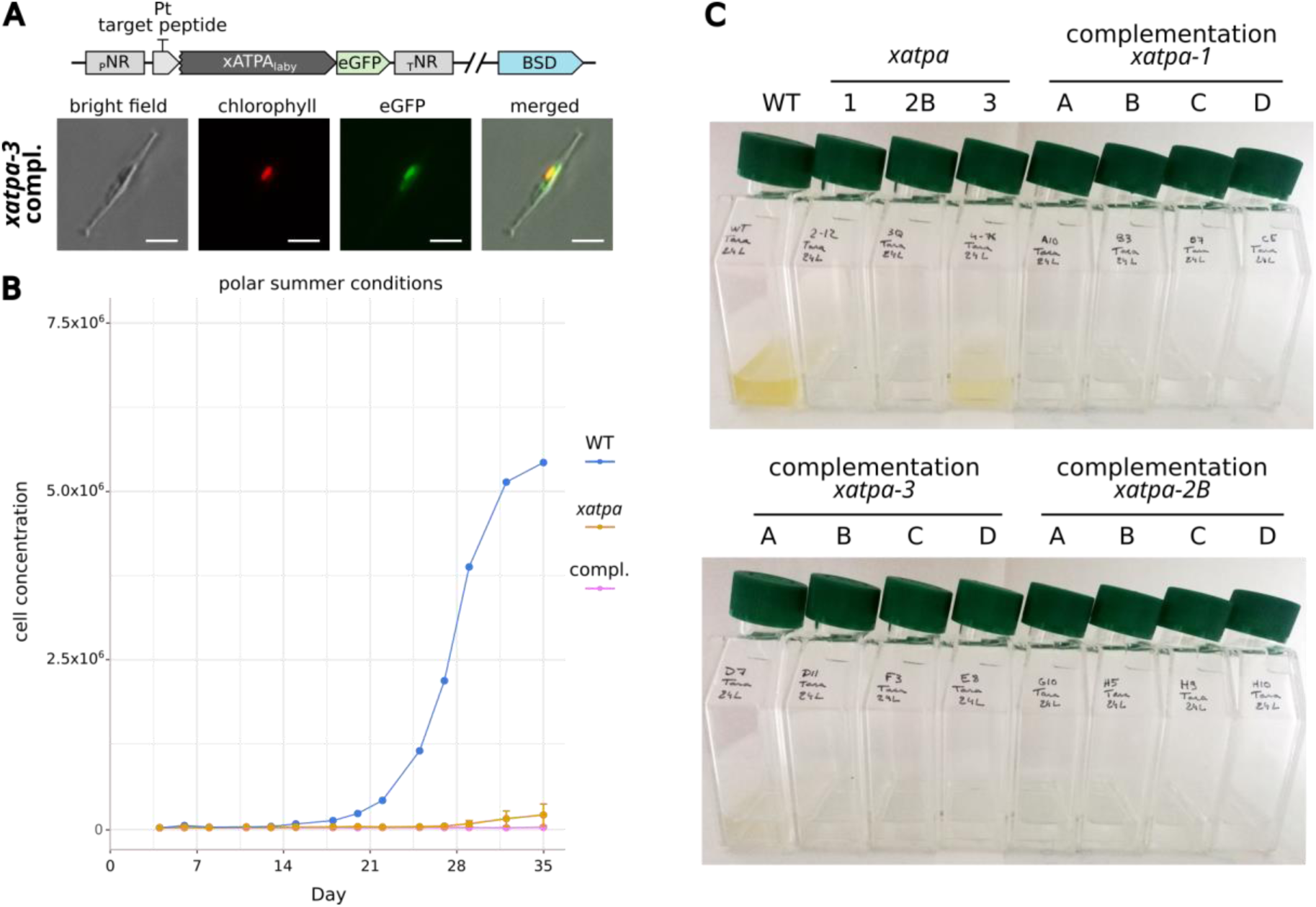
Complementation of *xatpa* strains with a labyrinthulomycete xATPA (*Aurantiochytrium sp.*, RCC893). A: A complemented *xatpa*-3 cells visualised by epifluorescence microscopy. The scale bars represent 5 μm. The genetic constructs are represented at the top of the image strip, showing the *Aurantiochytrium sp.* (RCC893) xATPA, whose targeting sequence is replaced by the *P. tricornutum* xATPA targeting sequence, fused with the eGFP. B: growth curve of WT (blue), *xatpa* (green) and complemented (orange) strains in polar summer conditions. Error bars represent the standard deviation of the mean. WT strain: n=1, *xatpa* strains: n=3, complemented strains: n=12. C: pictures of cultures at the end of the growth curve, showing very reduced growth of *xatpa* and complemented strains.

**Fig. S16.**
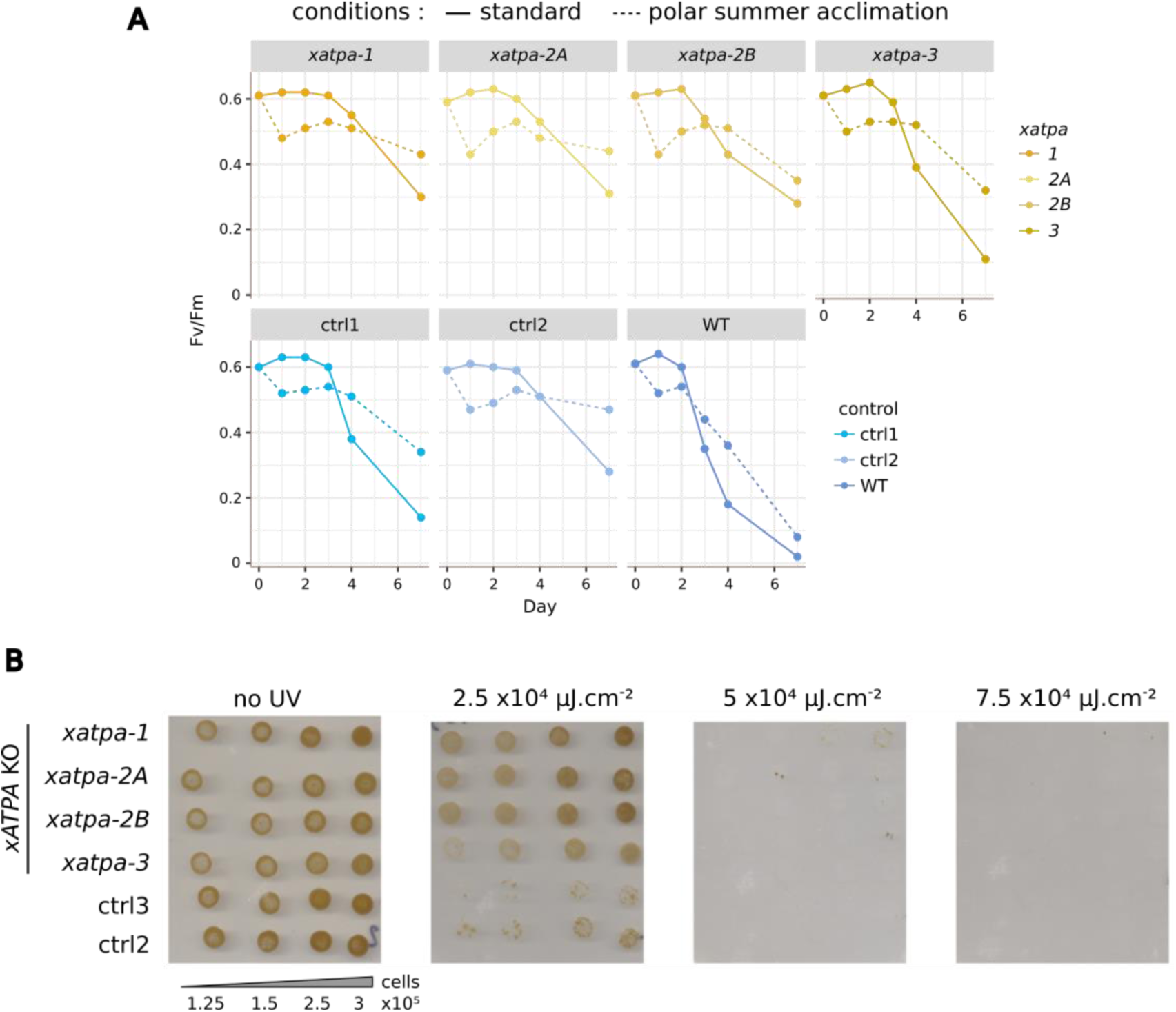
Light-related phenotypes of *xatpa* and control strains. A: Fv/Fm variation during culture in standard conditions or acclimation to polar summer conditions. Cultures in exponential phase were split in two (day 0) and either kept in standard conditions or transferred in polar summer conditions. Fv/Fm was measured for each strain over a week, at the day mid-point of the standard condition. Strong decreases in Fv/Fm values correspond to the end of the exponential phase. Note that a 3-day acclimation to polar summer conditions is used for RNAseq analysis, with *xatpa* and *ctrl* strains showing no alteration in their Fv/Fm. B: UV sensitivity of *xatpa* and *ctrl* strains was assessed on solid medium (35). Increasing number of cells from liquid cultures were spotted on solid medium and exposed to different intensities of UV light, and their survival was visually assessed after several days of recovery. UV intensity is indicated at the top of each pictures, lines correspond to strains indicated on the left, and columns corresponding to cell number per spot are indicated at the bottom.

**Fig. S17.**
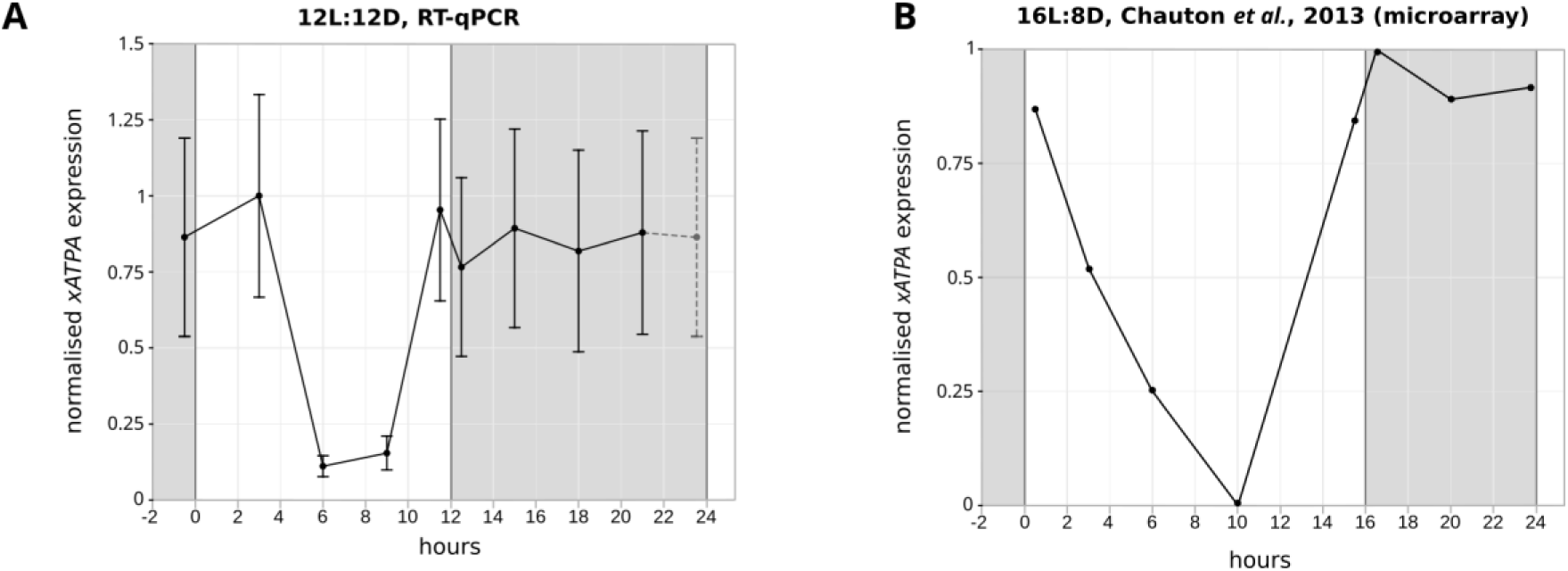
Diel dynamics of *xATPA* RNA levels in *P. tricornutum*. A: Cultures were grown in a 12h light: 12h dark (12L:12D) cycle, and *xATPA* RNA levels were assessed by RT-qPCR at different timepoints of the day. xATPA relative expression was calculated using the RPS and TBP housekeeping genes (36), and normalised by the highest mean value in each time series. Error bars corresponds to standard deviation of the normalised relative values (technical replicates, n=4). The light onset correspond to the 0 timepoint, grey areas represent the dark phases. The first time point is also reported as the last one (grey and dashed line) to give a clearer representation of the diel dynamic. B: xATPA (Phatr2_31465) dynamics from the microarray dataset published by Chauton et al., 2013 (46). These data were generated in a 18L:6D light regime, although xATPA exhibit a similar pattern as what is observed in 12L:12D. The light onset correspond to the 0 timepoint, grey areas represent the dark phases.

**Fig. S18.**
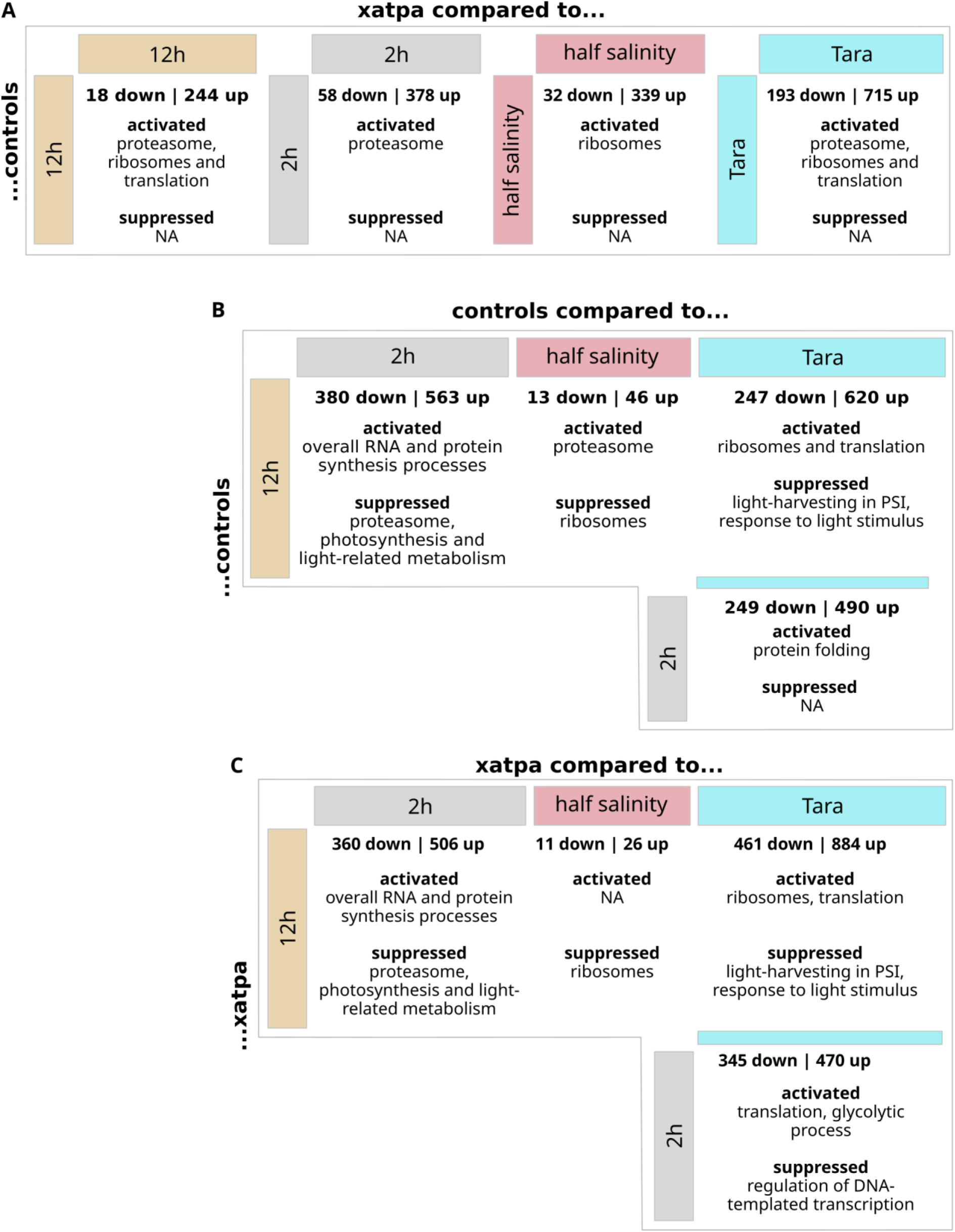
Summary of DEG and GSEA conducted across the whole dataset. For each pairwise comparison, the number of differentially down and upregulated genes (adjusted p-value < 0.05 and |log2FC| > 2) are indicated, as well as GO term enriched functions. Activated : significant positive enrichment of genes involved in a specific function, Suppressed : significant negative enrichment of genes involved in a specific function. A: Comparison between *xatpa* and *ctrl* strains in the same culture conditions. B: Comparison between *ctrl* strains in different culture conditions. C: Comparison between *xatpa* strains in different culture conditions.

**Fig. S19.**
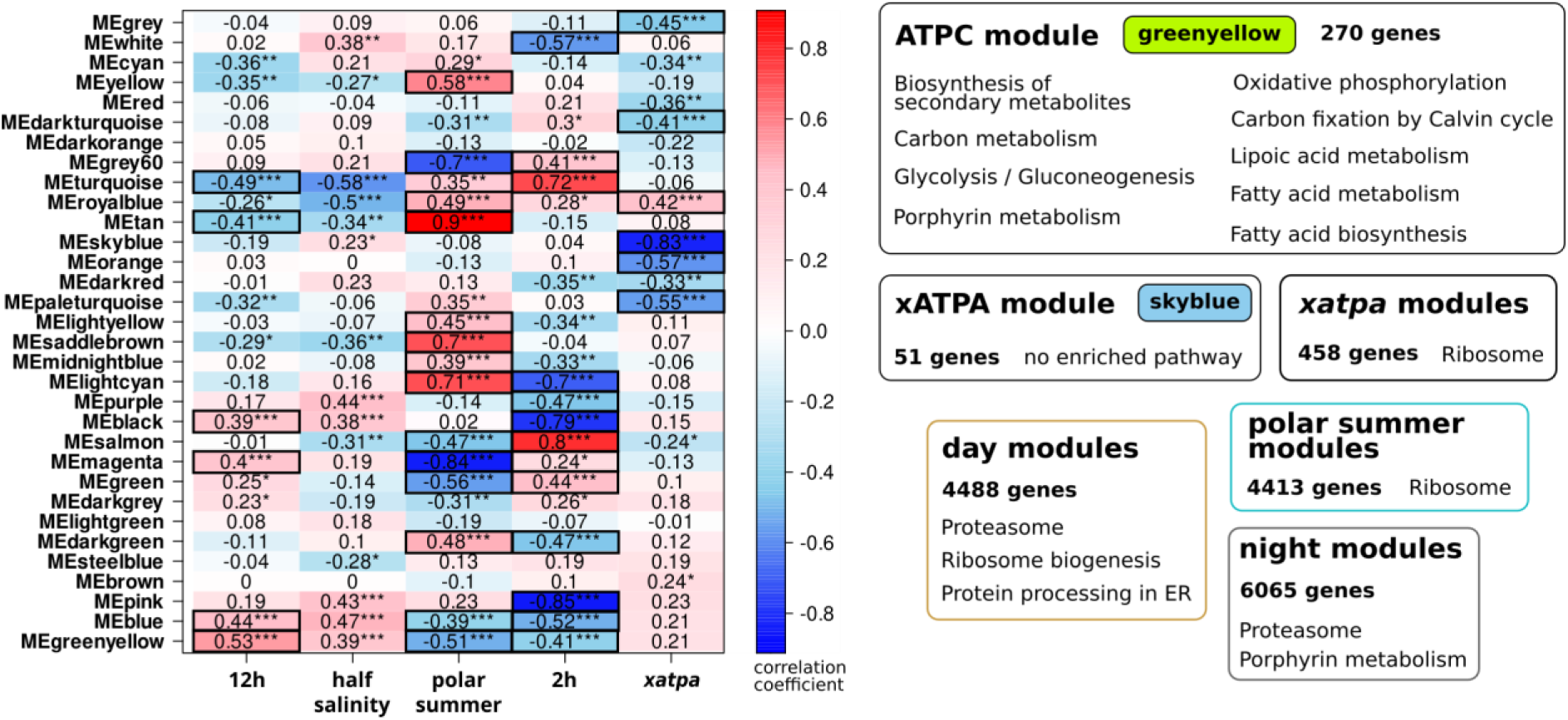
Weighted correlation network analysis (WGCNA) of the RNAseq dataset. Left: Correlation matrix between module eigeingenes and experimental conditions. The colour scale represents the Pearson correlation coefficient. Modules used in the functional enrichment analyses are indicated by a black box. Right: KEGG pathway enrichment of modules or groups of modules of interest. Enhancement or suppression of expression cannot be determined using this approach. Groups of modules were obtained by combining genes from modules with the most significant correlation coefficients (indicated by *** and surrounded by a black box), for each experimental condition. The number of genes considered in each analysis is shown in bold.

**Fig. S20.**
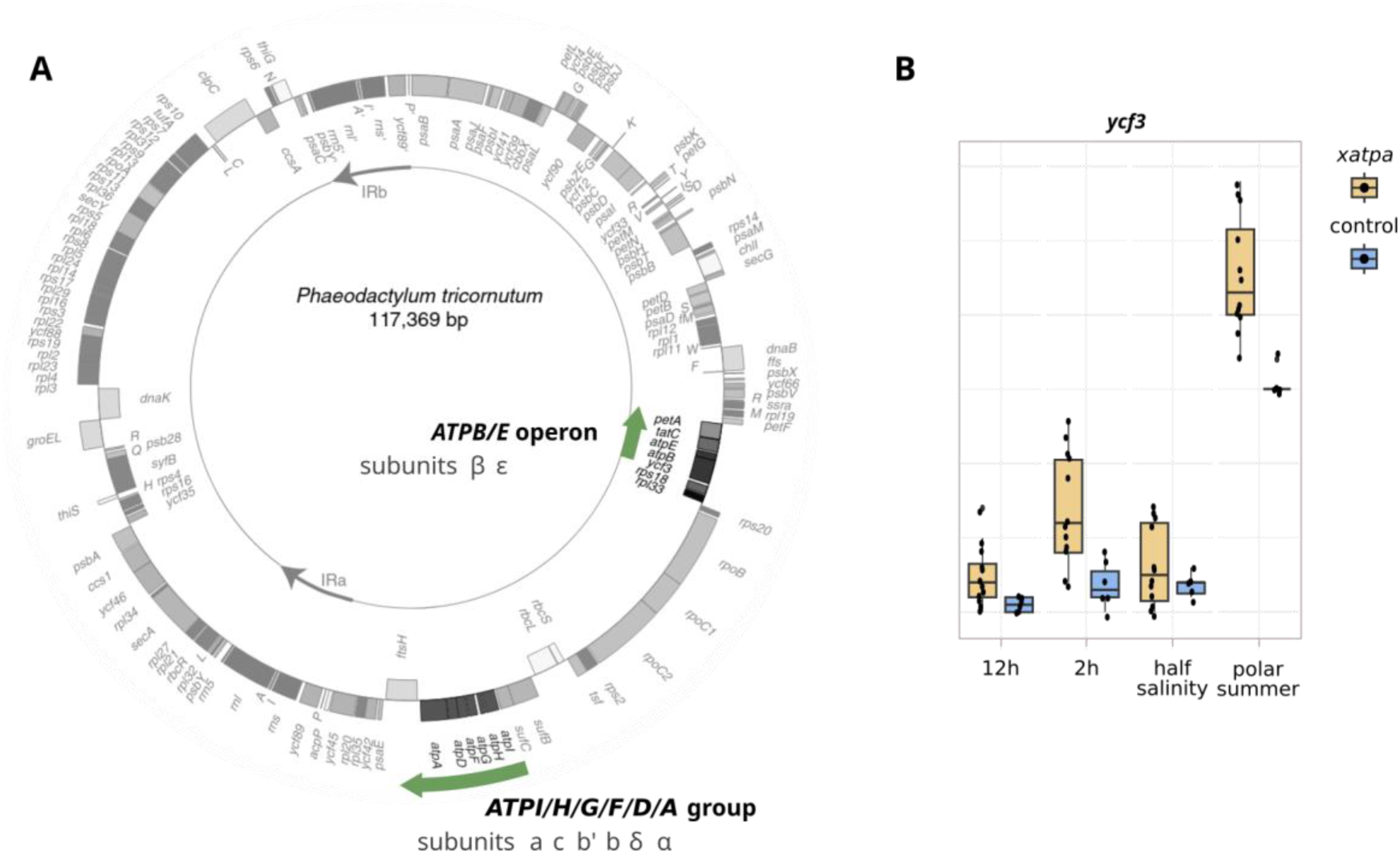
Rank analysis of chloroplast-encoded genes. A: Chloroplast chromosome of *P. tricornutum* showing the two *ATPB/E* and *ATPI/H/G/F/D/A* gene clusters. The corresponding subunit is indicated under each gene. Note the direct upstream position of *ycf3* from *ATPB*. Adapted from Oudot-Le Secq et al., 2007 (40). B: Comparison of rank distribution of the *ycf3* gene present on the 5’UTR of the *ATPB* gene, between *xatpa* and control strain, for individual growth conditions. Genes were ranked based on their gene counts, with a high rank value corresponding to a high gene count.

**Fig. S21.**
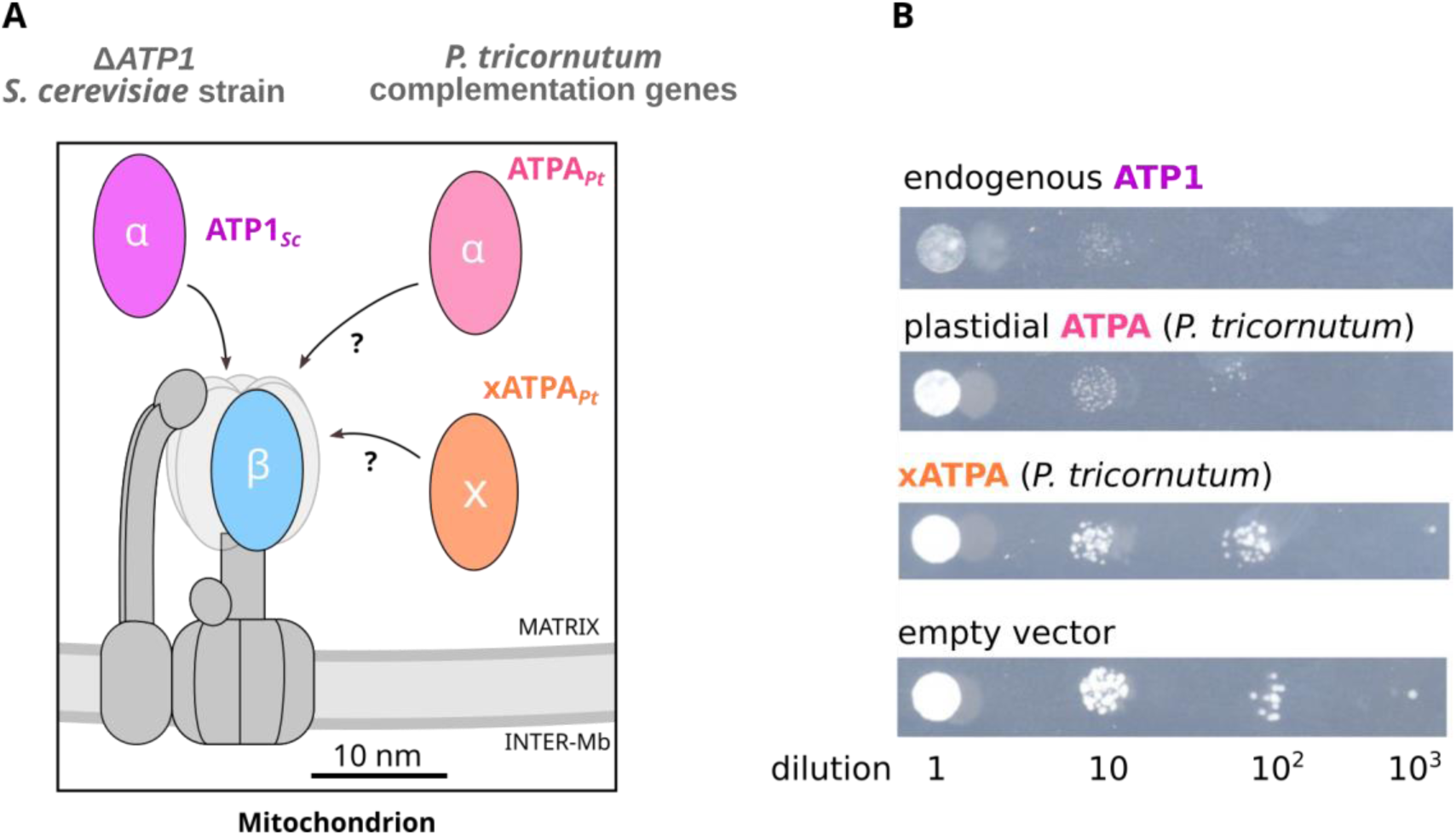
Effect of the heterologous overexpression of *P. tricornutum xATPA* and plastidial *ATPA* genes in the Δ*ATP1 Saccharomyces cerevisiae* yeast strain. A: BY4741 *S. cerevisiae* knocked-out for the nucleus-encoded mitochondrial α subunit (*ATP1*) was used as an *in vivo* model. Endogenous *ATP1*, *P. tricornutum* plastidial *ATPA* bearing the ATP1 mitochondrial targeting sequence (MTS) and *P. tricornutum xATPA* bearing the ATP1 MTS instead of its plastidial targeting sequence were overexpressed under the control of the strong GPD promoter. B: Spot assay on solid CSM-Ura 2% glucose medium. Exponential liquid cultures adjusted to DO=1 were spotted after serial dilutions. Pictures shown are representative of the 12 independent strains generated for each genetic construct. Δ*ATP1 Saccharomyces cerevisiae* strains overexpressing the endogenous ATP1 or ATPA*_Pt_* exhibit limited growth, likely due to cytotoxicity of accumulated α subunits. On the contrary, strains overexpressing xATPA*_Pt_* show similar growth abilites compared to the controls. This suggests different biological activities of xATPA compared to canonical α subunits *in vivo*. However, none of these strains could grow on CSM-Ura 2% glycerol as the original BY4741 strain, indicating that mitochondrial respiration could not be restored by canonical α subunit overexpression (cytotoxicity). Regarding xATPA*_Pt_*, this result suggests an absence of functional redundancy in the ATP synthase complex.

**Fig. S22.**
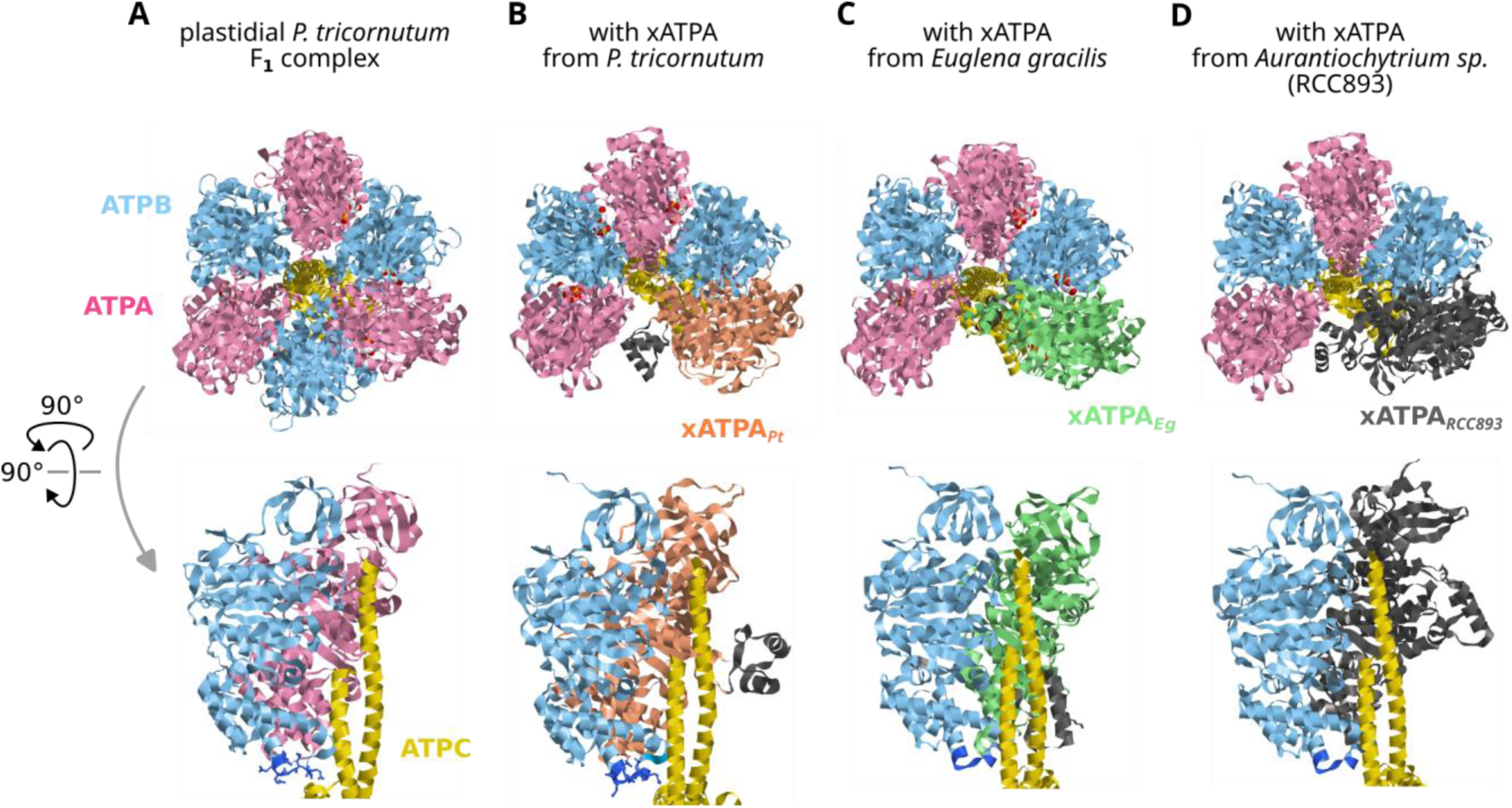
Predicted plastidial F_1_ complexes from *P. tricornutum* with various xATPA homologues. Predicted structures and interactions in F_1_ ATP synthase complexes using plastidial *P. tricornutum* ATPB (blue, UniProt: A0T0D2), ATPA (pink, UniProt: A0T0F1), ATPC (yellow, UniProt: Q84XB4). Proteins are shown as ribbons. Targeting sequences were removed from ATPC and xATPA for interaction computations. Top: Top view of the F_1_ complex. Bottom: Side view of the F_1_ complex, with only ATPC and two other subunits shown. The DELSEED amino acids of ATPB are represented in dark blue (D395-D401 amino acids). The xATPA bump domains are shown in grey. A: canonical 3α:3β:1γ complex (ipTM = 0.86, pTM = 0.87), B: complex including the *P. tricornutum* xATPA (orange, UniProt: B7FRE6) (ipTM = 0.77, pTM = 0.80). C: complex including the *Euglena gracilis* green-related xATPA (green) (ipTM = 0.74, pTM = 0.78). D: complex including the *Aurantiochytrium sp*. SAR-related xATPA (dark grey)(ipTM = 0.78, pTM = 0.81). Note that in F_1_ complexes with xATPA, the bump domain is predicted to consistently prevent the binding of a third β subunit, leading to an incomplete 1xATPA:2α:2β:1γ complex.

**Fig. S23.**
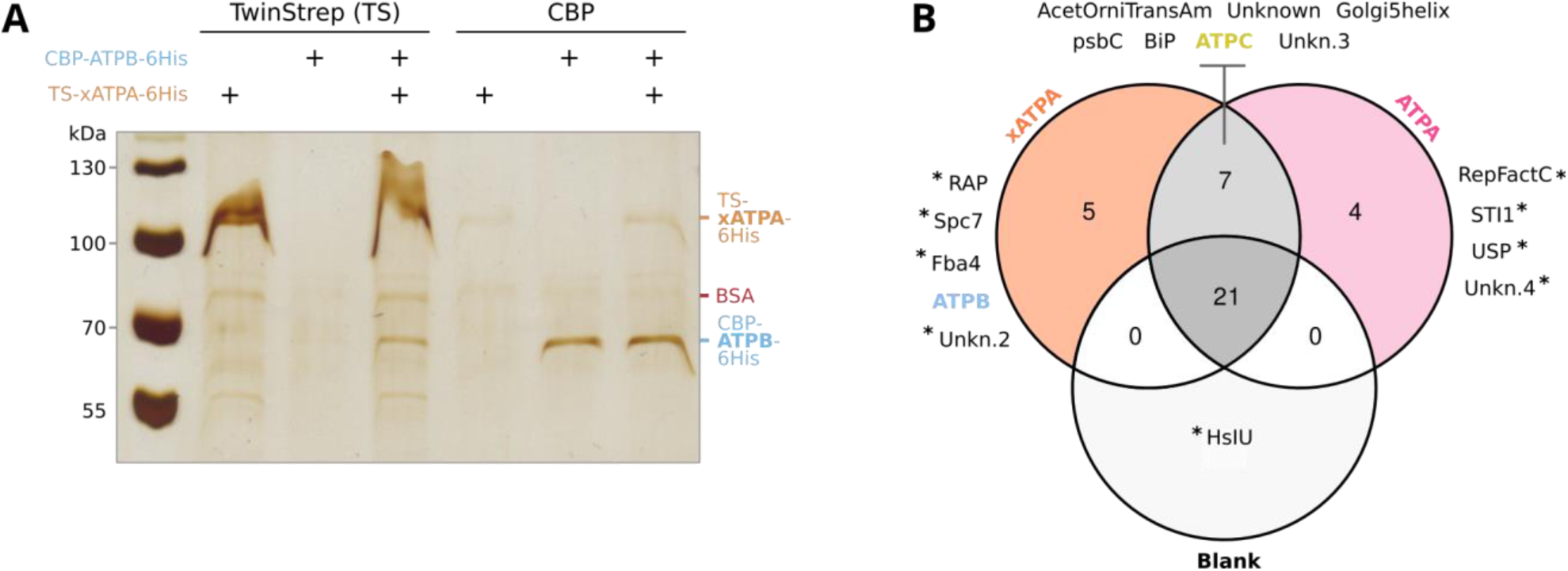
Interactions between *P. tricornutum* F_1_ subunits and xATPA*_Pt_*. A: Pull-down assays performed on TwinStrep-xATPA-6His and CBP-ATPB-6His in the presence of ATP and BSA. Input composition is shown on top of each lane, and the affinity tag used as bait is indicated above. Protein ladder sizes are indicated on the left, and band identities on the right. The gel was stained using silver nitrate. B: Putative protein partners identified from mass spectometry. Proteins were identified by mass spectrometry from pull-downs performed using whole cell lysates, using purified TwinStrep-xATPA-6His or TwinStrep-ATPA-6His as bait, or without tagged proteins (blank, TwinStrep affinity beads only). Protein baits are shown in bold and coloured. Short names of protein partners are displayed around the Venn diagram, with ATP synthase subunits shown in bold and coloured. Stars indicate proteins that are only detected once across all samples (n=3 for each sample type).

**Fig. S24.**
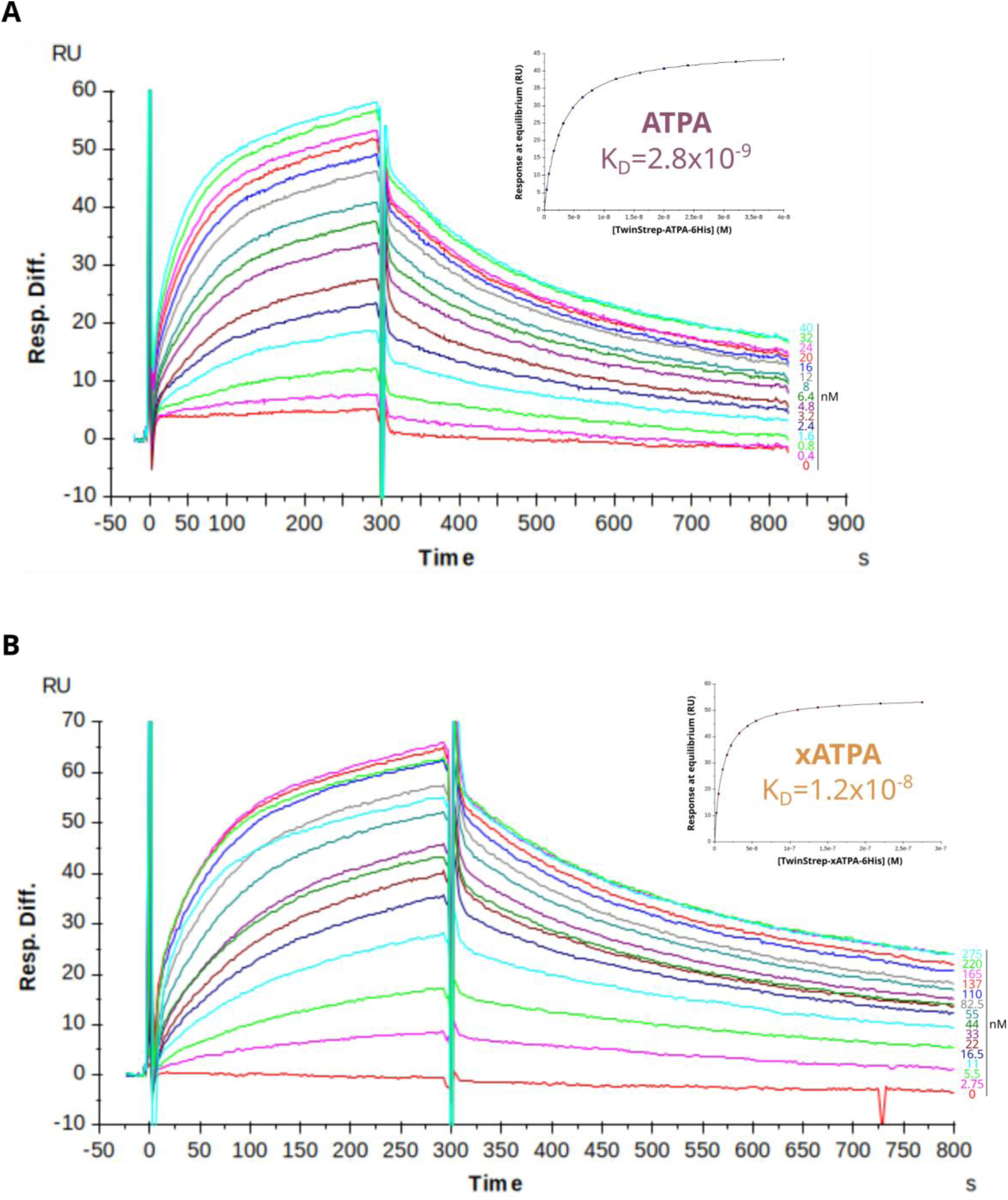
Comparison of ATPB affinity for ATPA and xATPA using Surface Plasmon Resonance. The association/dissociation kinetics between immobilised CBP-ATPB-6His and its TwinStrep-ATPA-6His and TwinStrep-xATPA-6His partners were measured via SPR. The same chip was subjected to increasing concentrations of recombinant ATPA and xATPA (colored curves), and interaction kinetics were computed. Top-right corner plots show the response at equilibrium (Req [RUs]) versus partner concentrations used for steady-state affinity fitting. Two technical replicates were performed on an independent chip to which ATPB was immobilised. The dissociation constant of the ATPB:ATPA dimer is 2.3x10^−9^±0.49 (n=2) while the ATPB:xATPA dimer is five times greater 1.2x10^−8^±0.1 (n=2), revealing a weaker affinity of xATPA for the β subunit than the canonical α subunit.

**Table S1.** UniProt ID of proteins shown in Fig 1. Other xATPA structures were predicted from their protein sequence with AlphaFold 3.

| Species | Protein | UniProt ID |
| --- | --- | --- |
| *Brassica oleracea* | ATPA (mito) | G4XYD7 |
| *Synechococcus elongatus* | ATPA | P08449 |
| *Lactobacillus acidophilus* | ATPA | Q5FKY2 |
| *Phaeodactylum tricornutum* | ATPA (cp) | A0T0F1 |
| *Phaeodactylum tricornutum* | xATPA | B7FRE6 |
| *Chlamydomonas reinhardtii* | xATPA | A0A2K3D914 |
| *Apatococcus fuscideae* | xATPA | A0AAW1T898 |
| *Ostreobium quekettii* | xATPA | A0A8S1IWP0 |
| *Guillardia theta* | xATPA | L1JSJ5 |
| *Seminavis robusta* | xATPA | A0A9N8E1V6 |

**Table S2.**
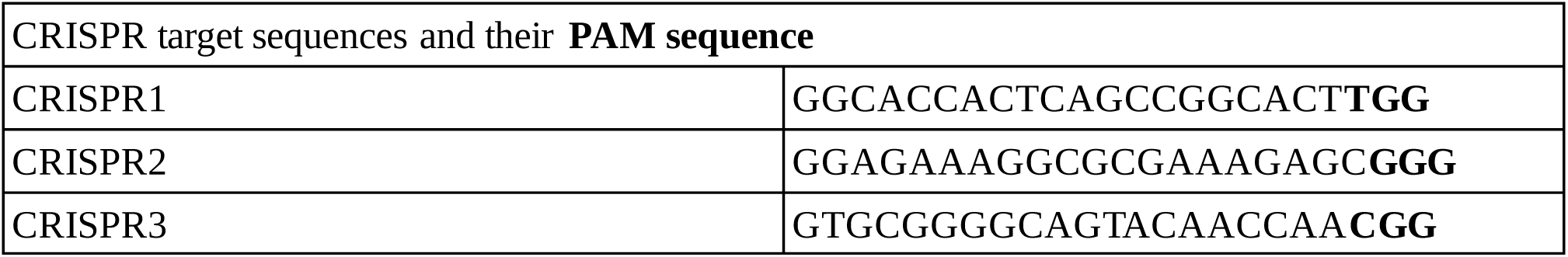
CRISPR sites successfully targeted on the *P. tricornutum xATPA* gene (Phatr3_J31465)

## Supplementary Data

**SI Dataset S1** (Dataset_S1.ods)

***Identification of xATPA homologues across the Tree of Life.*** Sheet 1: detailed information on genomes and transcriptomes used as a custom dataset. Sheet 2: reference ATPA sequences. Sheet 3: reference ATPB sequences. Sheet 3: list of the 222 xATPA sequences identified in this study. Sheet 4: first set of 27 xATPA sequences used as references for the subsequent identification of xATPA sequences.

**SI Dataset S2 (Dataset_S2.ods)**

***xATPA in the* Tara *Oceans dataset.*** Sheet 1: environmental xATPA sequences and their taxonimic classification; Sheet 2: environmental parameters and *xATPA* metagenomic abundances; Sheet 3: environmental parameters and *xATPA* metatranscriptomic abundances; Sheet 4: list and coordinates of *Tara* Oceans sampling stations; Sheet 5: normalisation data; Sheet 6: Random Forest dataset with train/test partition.

**SI Dataset S3 (Dataset_S3.ods)**

***List of oligonucleotides and plasmids sequences.*** Sheet 1: cloning and genotyping primers; Sheet 2: qPCR primers; Sheet 3: plasmid sequences.

**SI Dataset S4 (Dataset_S4.ods)**

*Composition of protein purification buffers.* Sheet 1: stock solutions; Sheet 2: buffers.

**SI Dataset S5 (Dataset_S5.ods)**

***RNAseq differential expression tables and mass spectrometry counts.*** Sheet 1: description; Sheet 2: MS/MS count values for all proteins identified in our samples; Sheet 3: custom gtf annotation file for the *P. tricornutum* chloroplast genome used to map reads of plastidial genes; Sheet 4: plastidial gene counts; Sheet 5: nuclear gene counts; Sheet 6 to 20: differential gene expression tables from pairwise comparisons.

## Notes

https://osf.io/89vm3/overview

### Competing Interest Statement

The authors have declared no competing interest.

